# The mechanochemical origins of the microtubule sliding motility within the kinesin-5 domain organization

**DOI:** 10.1101/2021.10.12.463902

**Authors:** Stanley Nithianantham, Malina K. Iwanski, Ignas Gaska, Himanshu Pandey, Tatyana Bodrug, Sayaka Inagaki, Jennifer Major, Gary J. Brouhard, Larissa Gheber, Steven S. Rosenfeld, Scott Forth, Adam G. Hendricks, Jawdat Al-Bassam

**Affiliations:** Dept. of Molecular and Cellular Biology, University of California, Davis, CA; Dept. of Biology, Dept. of Bioengineering, McGill University, Montreal, QC; Dept. of Biology, Rensselaer Polytechnic Institute, Troy; Dept. of Chemistry, The Ben Gurion University, Ber Sheva, Israel; Dept. of Pharmacology, Mayo Clinic, Jacksonville, FL

**Author notes:** These authors contributed equally to this work (Stanley Nithianantham, Malina K. Iwanski, Ignas Gaska). These authors are co-corresponding.

## Abstract

The conserved kinesin-5 bipolar tetrameric motors slide apart microtubules during mitotic spindle assembly and elongation. Kinesin-5 bipolar organization originates from its conserved tetrameric helical minifilament, which position the C-terminal tail domains of two subunits near the N-terminal motor domains of two anti-parallel subunits (Scholey et al, 2014). This unique tetrameric structure enables kinesin-5 to simultaneously engage two microtubules and transmit forces between them, and for multiple kinesin-5 motors to organize via tail to motor interactions during microtubule sliding (Bodrug et al, 2020). Here, we show how these two structural adaptations, the kinesin-5 tail-motor domain interactions and the length of the tetrameric minifilament, determine critical aspects of kinesin-5 motility and sliding mechanisms. An x-ray structure of the 34-nm kinesin-5 minifilament reveals how the dual dimeric N-terminal coiled-coils emerge from the tetrameric central bundle. Using this structure, we generated active bipolar mini-tetrameric motors from *Drosophila* and human orthologs, which are half the length of native kinesin-5. Using single-molecule motility assays, we show that kinesin-5 tail domains promote mini-tetramers static pauses that punctuate processive motility. During such pauses, kinesin-5 mini-tetramers form multi-motor clusters mediated via tail to motor domain cross-interactions. These clusters undergo slow and highly processive motility and accumulate at microtubule plus-ends. In contrast to native kinesin-5, mini-tetramers require tail domains to initiate microtubule crosslinking. Although mini-tetramers are highly strained in initially aligning microtubules, they slide microtubules more efficiently than native kinesin-5, due to their decreased minifilament flexibility. Our studies reveal that the conserved kinesin-5 motor-tail mediated clustering and the length of the tetrameric minifilament are key features for sliding motility and are critical in organizing microtubules during mitotic spindle assembly and elongation.

## Introduction

During cell division, microtubules (MTs) are rearranged into bipolar mitotic spindles from two overlapping MT asters during metaphase. Kinesin-5 motors are conserved across eukaryotes and are essential for the assembly and elongation of bipolar mitotic spindles. Kinesin-5 motors are 60-80 nm bipolar tetrameric proteins with a pair of N-terminal motor and C-terminal tail domains on either end of an α-helical central tetrameric minifilament (Kashina et al., 1996; Scholey et al., 2014). Although kinesin-5 motors are conserved across eukaryotes, there is a remarkable diversity in their mechanisms across species with the yeast kinesin-5 motors uniquely undergoing minus end directed motility as single motors, and switching direction upon clustering or during MT sliding toward plus-end directed motility (Pandey et al., 2021a; Pandey et al., 2021b; Shapira et al., 2017; Singh et al., 2018). In contrast, metazoan Kinesin-5 motors undergo motility towards MT plus ends both along single MTs and when sliding MTs (Bodrug et al., 2020; Kapitein et al., 2005; Weinger et al., 2011).

Kinesin-5 motors slide apart MTs emanating from opposing spindle poles at the midzone region by initially crosslinking, then aligning these MTs by undergoing motility with their bipolar ends along two MTs using their unique tetrameric architecture (Kashina et al., 1996). This kinesin-5 MT sliding activity is conserved and critical for organizing bipolar mitotic spindles. The kinesin-5 tetrameric bipolar organization originates from the anti-parallel folding of the four α-helices from the four subunits within the Bipolar assembly (BASS) domain (Scholey et al., 2014). The BASS domain lies at the center of the kinesin-5 minifilament and forms the central force-bearing structure that coordinates between the two motile motor ends, each of which supports hand over-hand motility along MTs. While the kinesin-5 MT sliding motility is linked to the motor’s bipolar tetrameric organization, the functional relationship between the conserved kinesin-5 motor, tail or BASS domains and the unique MT crosslinking or MT sliding mechanisms remains unknown (Kapitein et al., 2005).

The kinesin-5 C-terminal tail domains (termed tail from herein) emerge from bipolar tetrameric minifilament near the motor domains of the antiparallel dimeric folded subunits (Acar et al., 2013) and kinesin-5 tails are essential for MT sliding activity across species (Duselder et al., 2015; Hildebrandt et al., 2006; Weinger et al., 2011). We recently discovered that the kinesin-5 tail regulates the MT-activated ATP hydrolysis in the motor domains and this regulation is essential for kinesin-5 motors to transition from crosslinking to MT sliding motility (Bodrug et al., 2020). This allosteric tail to motor domain interaction allow for trans interactions between multiple kinesin-5 motors, resulting in multi-motor clustering (Bodrug et al., 2020). Tail-mediated clustering is a critical for organizing the forces generated by multiple kinesin-5 motors promote the alignment of MTs to crosslinked paired MTs then modulate their sliding (Bodrug et al., 2020). However, it remains unknown how this tail-to motor regulation is impacted by the distinct kinesin-5 central minifilament organization and how these two features regulate MT crosslinking, alignment and sliding.

Here, we describe how the kinesin-5 tail-motor domain interaction and the length of the tetrameric minifilament modulate kinesin-5 motor clustering, MT crosslinking, and MT sliding activities. We determined an x-ray structure of a 34-nm extended BASS α-helical tetramer, revealing rigid dimeric parallel N-terminal coiled-coil junctions that emerge from its central tetrameric core. Using this structure as a platform, we engineered short human and *Drosophila* bipolar 38-nm mini-tetrameric kinesin-5 motors, which are roughly half the length of native kinesin-5 motors. Single-molecule motility assays reveal mini-tetramers without tail domains undergo processive motility with infrequent pauses in which motors statically bind MTs without diffusing. The tail domains enhance kinesin-5 mini-tetramer pausing and promote the assembly of multiple motors into clusters. These multi-motor clusters assemble when motile motors encounter paused motors along the MT. MT sliding assays reveal that unlike the native kinesin-5 motor, kinesin-5 mini-tetramers require the tail domain to crosslink, align and slide two MTs. Kinesin-5 mini-tetramers are restricted in their ability to pair and align MTs. However, once MTs are paired, these mini-tetramers slide these MTs more efficiently than native kinesin-5. Our data demonstrate that the kinesin-5 tail and length of the minifilament are two critical structural adaptations that allow it to effectively crosslink and slide apart MTs. The tail domain is required to mediate motor pausing and clustering such that multiple motors can coordinate in MT overlaps. The length of the kinesin-5 minifilament is critical to provide flexibility required for efficient MT crosslinking and transmission of forces during MT alignment and sliding.

## Results

### An extended BASS x-ray structure reveals rigid and dimeric coiled-coils emerging from bipolar tetramer junctions

To determine the organization of bipolar kinesin-5 minifilament and how the dimeric α-helical coiled-coil junctions are formed, we solved the x-ray structure of an extended *Drosophila melanogaster* KLP61F segment (residues 620-804) in which the kinesin-5 sequence is extended by 40 residues N-terminally compared to the previous BASS x-ray structure (Scholey et al 2014; termed BASS-XL from herein). The KLP61F BASS-XL was purified and crystallized in the space group C2 (253.18-Å, 84.89-Å, 96.77-Å) (Figure 1 figure supplement 1). The X-ray diffraction data were highly anisotropic, indicating translational pseudo-symmetry, and were thus elliptically truncated to 4.4-Å resolution in reciprocal space axes (Figure 1 figure supplement 1; Table 1). The BASS-XL structure was determined by molecular replacement using BASS structure as a starting model and was refined to 4.4-Å resolution leading to a R_work_/R_free_ (0.277/0.309). The 4.4-Å BASS-XL x-ray crystal structure reveals a 34-nm α-helical minifilament, compared to the 27-nm BASS minifilament (Figure 1, figure supplement 1A). The structure reveals the N-terminal 30-residues form parallel coiled-coils which emerge from both ends of the BASS tetrameric core (Figure 1A)(Scholey et al., 2014). The N-terminal parallel coiled-coils form multiple heptad repeat interactions with clear a and d contacts forming homotypic interfaces (residues 620-670) (Figure 1B). These dimeric α-helical coiled-coil extend for 10-nm before they twist slightly out of register into a swap junction (residues 693-697) to form a four α-helical anti-parallel bundle within the BASS core tetramer (residues 697-760)(Figure 1A)(Scholey et al., 2014). The C-terminus of the BASS-XL forms α-helices (residues 760-806), which stabilizes a junction of the N-terminal coiled-coil dimer at each end of the BASS tetramer core for the subunit that emerges from the opposite end (Figure 1A). This junction rigidly and tightly orients the BASS-XL N-terminal coiled-coils onto the ends of the BASS core and positions the N-terminal ends of the coiled-coil to be 180° with respect to those emerging from the opposite end of the BASS-XL structure (Figure 1A).

**Figure 1:**
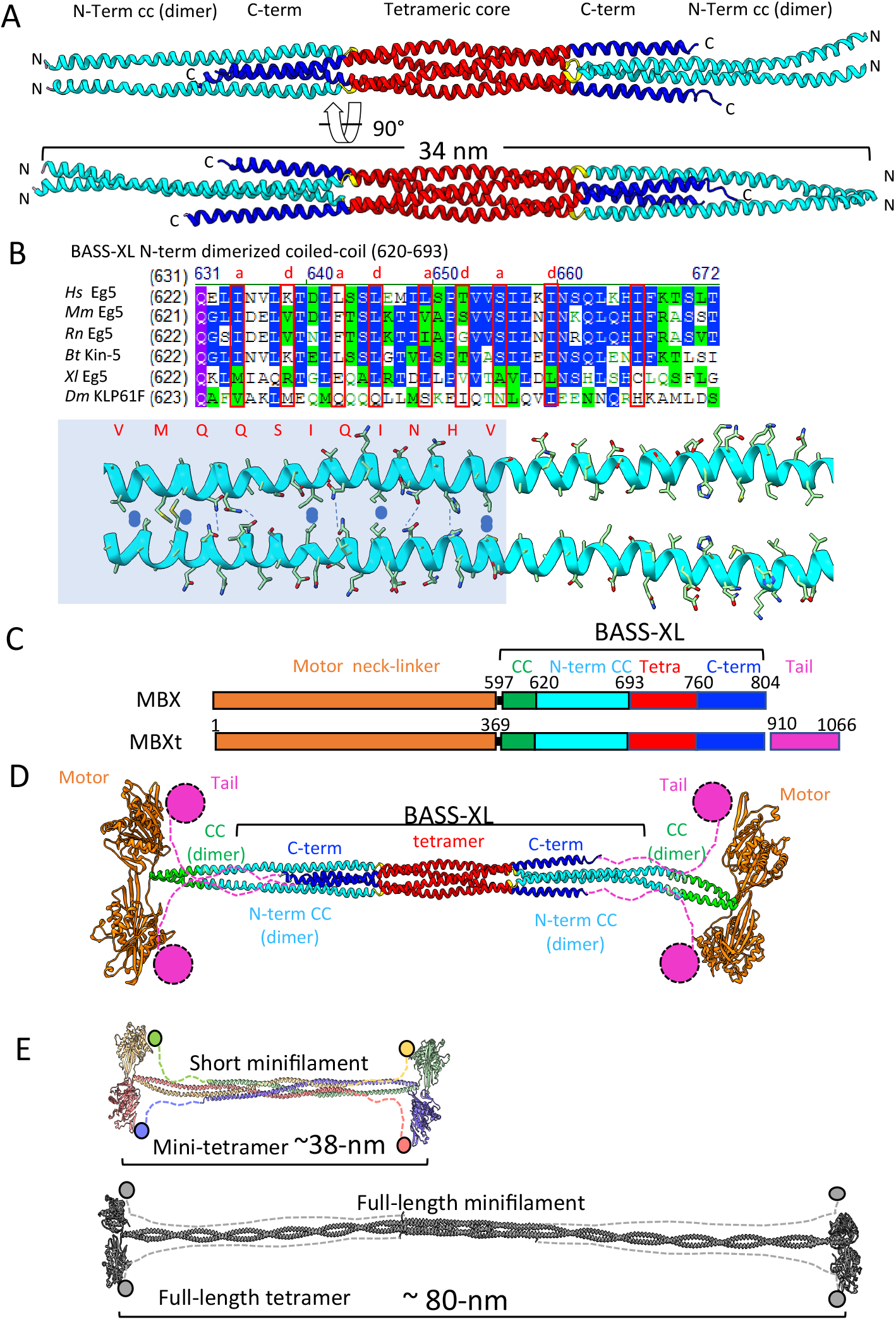
The x-ray structure of the kinesin-5 BASS-XL minifilament reveals rigid dimeric coiled-coils emerge from a tetramer core and allow designing kinesin-5 mini-tetramers. A) Top, a side views of 34 nm long KLP61F BASS-XL minifilament crystal structure reveals the formation of dimeric coiled-coils (cyan: 40-nm long) that are rigidly attached to the BASS tetrameric core (red) and stabilized by the C-terminal extension (dark Blue). Bottom panel, 90°rotated view compared to top panel. B) Top sequence alignment reveals the conservation, and the heptad repeat pattern of the dimeric coiled-coil (marked in a and d residues), mapped based on the x-ray BASS-XL structure. Bottom, view of the dimeric coiled-coil heptad interactions marked in B on the structural interface shown in A C) Domain organization of the designer Kinesin-5 mini-tetramers. The Motor and neck linker (residues 1-369; Blue) are fused to 20 residue N-terminal coiled-coils (green) and then fused to the N-terminal dimeric-coiled coil junctions (cyan) of the BASS-XL and the tetramer core (red) followed by the C-terminal extension (deep blue). The mini-tetramers may either include or lack the Kinesin-5 C-terminal tail domain (orange). D) The structural organization of the designed 38-nm long kinesin-5 mini-tetrameric motor based on the fusion of domains based on matching the heptad repeats with neck helical coiled-coil. The tail domains extend near the motor domains of the subunits folded in anti-parallel orientations. E) Scaled-comparison of the mini-tetramer kinesin-5 motors (38 nm), shown on the top, reveals that they are half the length of the native kinesin-5 tetrameric motors (80 nm), shown on the bottom.

**Table 1.** X-ray crystallographic Data collection and refinement statistics.

|  | BASS-XL |
| --- | --- |
| **Data collection** |  |
| Resolution range | 126.55- 4.40 (4.64 - 4.40) |
| Wavelength (Å) | 0.979 |
| Space group | C 2 |
| Unit cell (Å): *a*, *b*, *c* | 253.18, 84.89, 96.77 |
| (°): $\beta$ | 91.44 |
| Total number of observed reflections | 55597 (7964) |
| Unique reflections | 13060 {10508}† |
| Average mosaicity | 0.33 |
| Multiplicity | 4.3 (4.3) |
| Completeness (%) | 98.7 (98.8) {79.76}† |
| Mean $\langle I \rangle / \sigma(I)$ | 4.6 (1.8) |
| Wilson B-factor | 38.14 |
| $R_{\text{merge}}^a$ | 0.10 (0.47) |
| **Structure refinement** |  |
| $R_{\text{work}}$ | 0.274 (0.251) |
| $R_{\text{free}}$ | 0.310 (0.368) |
| Molecules per asymmetric unit | 4 |
| Number of non-hydrogen atoms | 5037 |
| macromolecules | 5037 |
| Solvent | 0 |
| Protein residues | 702 |
| RMS bond lengths (Å) | 0.002 |
| RMS bond angles (°) | 0.56 |
| Ramachandran favored (%) | 96.2 |
| Ramachandran allowed (%) | 3.6 |
| Ramachandran outliers (%) | 0.43 |
| Rotamer outliers (%) | 0.41 |
| Clashscore | 11.42 |
| Mean *B* values (Å<sup>2</sup>) |  |
| Overall | 113.93 |
| macromolecules | 113.93 |
| Number of TLS groups | 11 |

Parenteses numbers represent the highest-resolution shell.
†Numbers represent the truncated data after treated with ellipsoidal truncation and anisotropic scaling.

Using the BASS-XL x-ray structure as template for a shortened kinesin-5 minifilament, we engineered bipolar kinesin-5 motors with a 38-nm central minifilament, which we term kinesin-5 mini-tetramers from herein (Figure 1C). Using the pattern of heptad repeats observed in the BASS-XL structure, a longer KLP61F BASS-XL sequence was utilized to generate mini-tetrameric motors by including three coiled-coil heptad repeats (21 residues) on the coiled-coil N-terminus observed in the BASS-XL x-ray structure, leading to a 38-nm long kinesin-5 minifilament (Figure 1 green; 21 residues 597-620). The N-terminal end of this revised 38 nm minifilament was then fused to the C-terminal end of the kinesin-5 motor domain and neck linker (residues 1-365) sequences. On its C-terminus, the BASS-XL mimi-tetramer sequence was either terminated or was C-terminally fused to the N-terminal end of the kinesin-5 tail domain (residues 903-1033) (Figure 1C; Figure 1 figure supplement 2). We generated a structural model for the 38-nm kinesin-5 mini-tetramer, showing its short overall length, which are roughly half the length of the native kinesin-5 (38 nm versus 80 nm)(Acar et al., 2013; Scholey et al., 2014) (Figure 1 figure supplement 2C-D; Figure 1D). The kinesin-5 mini-tetramer model shows that BASS-XL rigidly orients the two dimeric motor domain pairs on both of its ends, positioning them 180° rotated with respect to each other (Figure 1D; Figure 1 figure supplement 2C-D). Using this strategy, we generated two mini-tetrameric constructs from two kinesin-5 orthologs and compared their activities: *Drosophila* KLP61F mini-tetramers (-tail: kMBX, or + tail: kMBXt) or human Eg5 kinesin-5 mini-tetramer motors (-tail: hMBX, or +tail: hMBXt) as described (Figure 1 figure supplement 1A-C; Figure 1C). The motor to coiled-coil junctions for the mini-tetramer constructs are predicted to lack flexibility with the respect to BASS tetrameric core, and to exhibit less torsional flexibility than the full-length native kinesin-5 (Figure 1D). The kinesin-5 mini-tetramer constructs represent only 60% of the full-length kinesin-5 sequence, as they lack 280 residues coiled-coil region between the motor neck linker and the extended N-terminal end of the BASS-XL structure, and 100 residues flexible region between the BASS-XL C-terminal domain and the N-terminal end of the tail domain (Acar et al., 2013; Bodrug et al., 2020; Scholey et al., 2014).

### Kinesin-5 mini-tetramers undergo processive plus end-directed motility along with pausing

We reconstituted the motility of kMBX and kMBXt mini-tetramers fused to Neon Green (mNG) along single MTs imaged them using total internal reflection fluorescence (TIRF) microscopy, as previously described (Bodrug et al., 2020). Single kMBX mini-tetramer motors undergo slow processive motility along MTs interspersed very seldom with brief pauses (Figure 2A-B). In contrast, kMBXt motors undergo processive motility towards MT plus-ends and pause for extended periods (Figure 2C-D). While paused along MTs, kMBXt motors often encounter other motile motors and merge to form brighter multi-motor clusters. These clusters of kMBXt motors undergo motility together as singular entities until reaching MT plus-ends, where they concentrate for extensive time periods. Thus, kMBXt motors pause for extended periods and form multi-motor clusters, which are not observed in the case of the kMBX motors, suggesting that the kinesin-5 tail-motor domain interaction promotes pausing and multi-motor clustering, recapitulating activities observed for full-length Eg5 (Bodrug et al., 2020).

**Figure 2:**
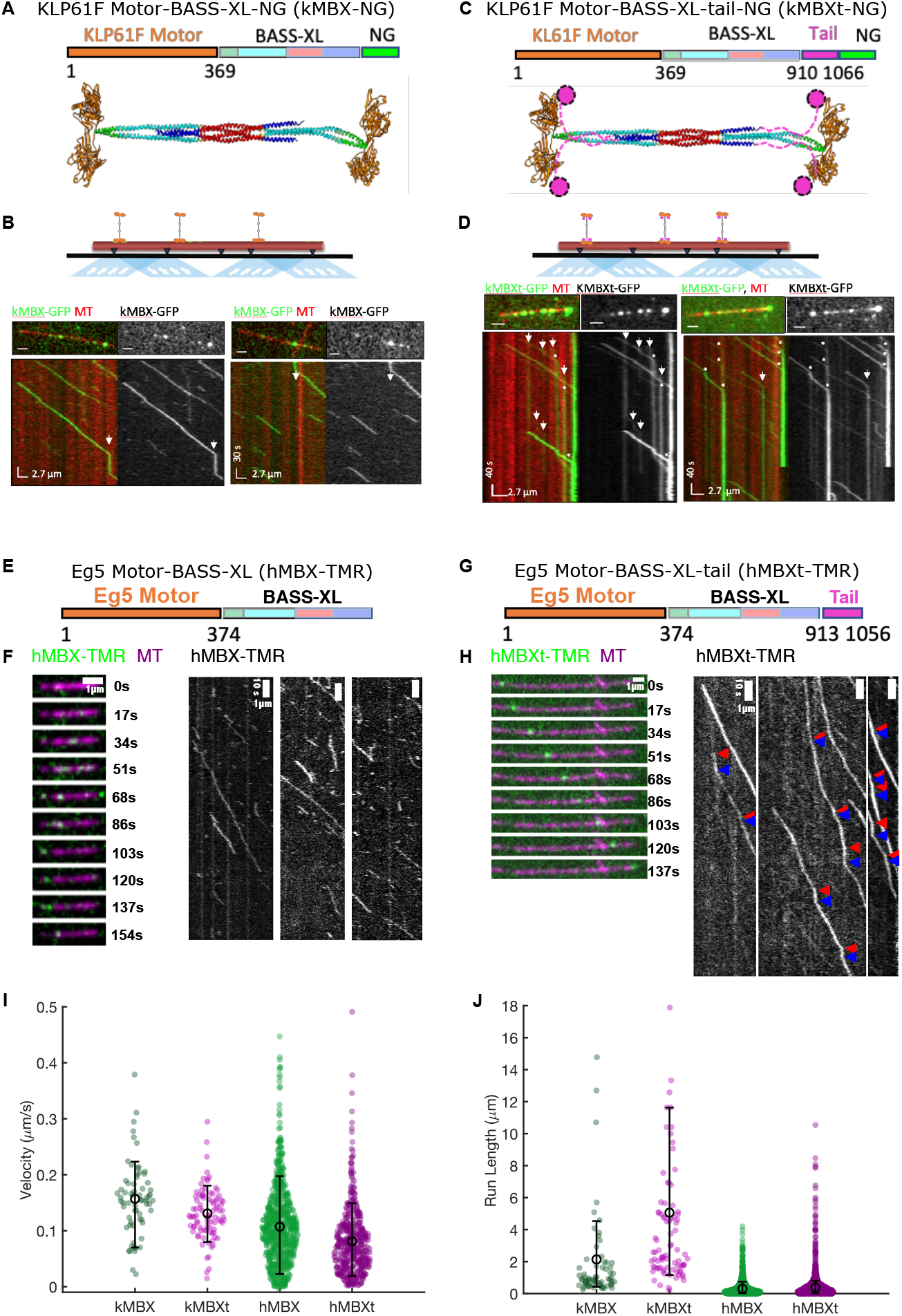
Kinesin-5 mini-tetramers undergo processive motility interrupted by static pauses along MTs *in vitro*. A) Top, kMBX domain organization. Motor and neck linker domains (1-365, blue) extended coiled-coil (green), BASS-XL minifilament with its dimerized zone (orange), tetrameric zone (red) and C-terminal zone (cyan). Middle, structural model for the kMBX mini-tetramer as shown in Figure 1D. B) Top, TIRF microscopy reconstitution setup to examine kMBX motors motility along MTs. Middle panel, Image of individual MT (red) with kMBX motors moving along toward Mt plus ends (right side). Bottom panel, Kymograph of above image with kMBX motor motility along MTs revealing their processive motility with extended pauses (arrows) and lack of accumulation at MT plus ends. C) Top, kMBX domain organization. Motor and neck linker domains (1-365, orange) extended coiled-coil (green), BASS-XL minifilament with its dimerized zone (cyan), tetrameric zone(red), C-terminal zone (blue) and C-terminal tail domain (pink). Middle, structural model for the kMBX mini-tetramer as shown in Figure 1D. D) Top, TIRF microscopy setup for kMBXt MT motility assays. Middle panel, Images of individual MTs (red) with KMBXt motors (green) moving along with MT plus end to the right. Bottom panel, kymograph of above image with KMBXt motor motility along MTs revealing their extended pausing and clustering (marked by arrows) and accumulating kMBXt motor at MT plus ends. E) The hMBX-TMR mini-tetramer consists of the human Eg5 motor-neck linker domain (1 374, orange), the Drosophila BASS domain (597-799) with its extended coiled-coil (green), with its dimerized zone (cyan), tetrameric zone (red), C-terminal zone (blue). F) Kymographs of hMBX mini-tetramer motor show that undergo processive motility towards MT plus-ends and pause infrequently. G) The hMBXt-TMR mini-tetramer consists of the human Eg5 motor-neck linker domain (1374, orange), the *Drosophila* BASS-XL minifilament (597-799) with its extended coiled-coil (green), with its dimerized zone (cyan), tetrameric zone(red), C-terminal zone (blue) and the human Eg5 C-terminal tail domain (913-1056, pink). H) An example of a very long run observed for hMBXt motors are shown. 20 frames between time points. Kymographs for motors showing the presence of pauses between periods of processive motility. The start (red) and end (blue) points of pauses are identified. I) Histogram distributions for motility velocity (μm/s) of the kMBX (green), kMBXt (pink), hMBx (green) and hMBXt (pink) motors along MTs showing that the kMBXt or hMBXt undergo slower motility than the kMBX and hMBX motors. J) Histogram distributions for motility run lengths (μm) the kMBX (green), kMBXt (pink), hMBx (green) and hMBXt (pink) motors along MTs revealing that the kMBXt or hMBXt motors are generally more processive than the hMBX and kMBX motors.

To understand how the *Drosophila* KLP61F tail domain regulates the mini-tetramer motor properties, we studied how motility of the kMBX and kMBXt are affected by changes in ionic strength (25, 50, 100 mM KCl). At 25 mM KCl, kMBXt motors undergo very slow motility, which is 30% slower than kMBX motors (Table 2: 78 vs 115 nm/s). At 50 mM KCl, both kMBX and kMBXt undergo slightly faster motility, with velocity of kMBX motor being about 10% faster than kMBXt motors (Table 2: 127 nm/s vs 144 nm/s). At 100 mM KCl, the average motility velocity of kMBX is 45% higher compared to that for kMBXt motors (Table 2:156 nm/s vs 114 nm/s). At higher ionic strengths, kMBXt velocities progressively decreased (Table 2). In contrast, the kMBX motors bound and dissociated rapidly from MTs at the higher salt concentrations and did not undergo processive motility. At 25 mM KCl, kMBX motors undergo extremely long run lengths. At 50 mM KCl, kMBX motors undergo motility at 66% shorter average run lengths than the kMBXt motors (Table 2:1014 vs 3135 nm) and 66% shorter run time (Table 2:9 vs 29 s). At 100 mM KCl, the kMBX motor undergo motility with a 60% shorter average run length compared to KMBXt (Table 2: 2137 nm vs 5060 nm), and 60% shorter average run time of kMBXt (Table 2:16 s vs 51 s). These data suggest that the kinesin-5 tail domain down-regulates the motor domain by decreasing its MT activated ATPase leading to slower motility but increasing run lengths and run times during each processive motility event.

**Table 2:**
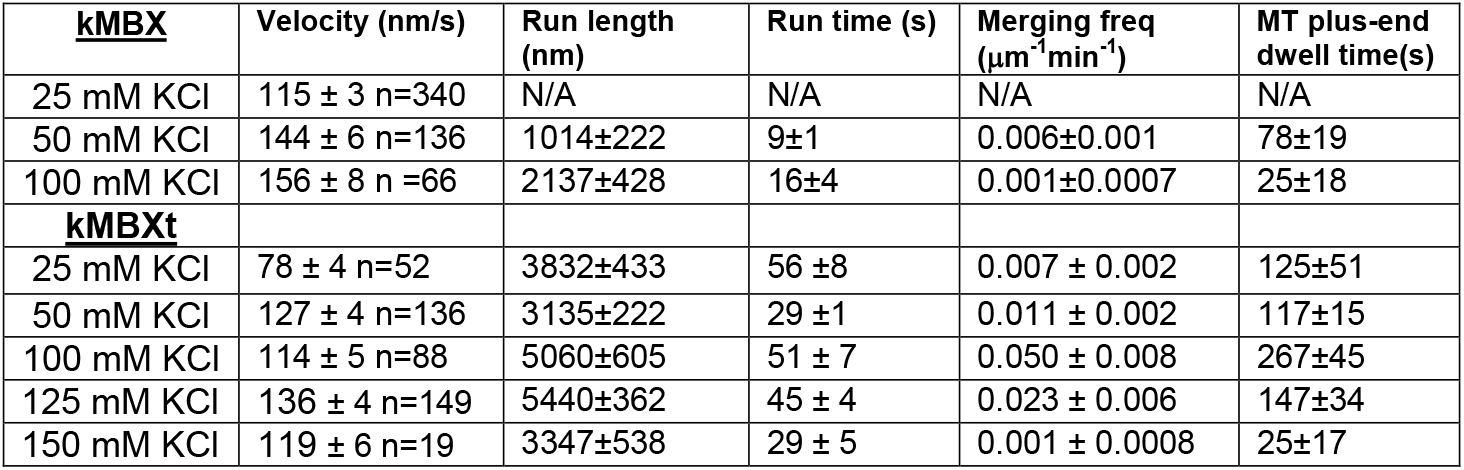
*Drosophila* Kinesin-5 mini-tetramer motor motility properties

| **<u>kMBX</u>** | **Velocity (nm/s)** | **Run length (nm)** | **Run time (s)** | **Merging freq ($\mu\text{m}^{-1}\text{min}^{-1}$)** | **MT plus-end dwell time(s)** |
| --- | --- | --- | --- | --- | --- |
| 25 mM KCl | 115 ± 3 n=340 | N/A | N/A | N/A | N/A |
| 50 mM KCl | 144 ± 6 n=136 | 1014±222 | 9±1 | 0.006±0.001 | 78±19 |
| 100 mM KCl | 156 ± 8 n =66 | 2137±428 | 16±4 | 0.001±0.0007 | 25±18 |
| **<u>kMBXt</u>** |  |  |  |  |  |
| 25 mM KCl | 78 ± 4 n=52 | 3832±433 | 56 ±8 | 0.007 ± 0.002 | 125±51 |
| 50 mM KCl | 127 ± 4 n=136 | 3135±222 | 29 ±1 | 0.011 ± 0.002 | 117±15 |
| 100 mM KCl | 114 ± 5 n=88 | 5060±605 | 51 ± 7 | 0.050 ± 0.008 | 267±45 |
| 125 mM KCl | 136 ± 4 n=149 | 5440±362 | 45 ± 4 | 0.023 ± 0.006 | 147±34 |
| 150 mM KCl | 119 ± 6 n=19 | 3347±538 | 29 ± 5 | 0.001 ± 0.0008 | 25±17 |

To explore whether the tail-motor interaction and their regulatory mechanism observed for *Drosophila* KLP61F are conserved in human Eg5, we generated human Eg5 motor minitetramers hMBX (-tail) and hMBXt (+tail) (Figure 2 2E-H). The hMBX and hMBXt motors were engineered with reactive cysteines for labeling with Tetra-methyl Rhodamine (TMR) fluorophores (see materials and methods). The bright, stable TMR fluorophores on hMBX and hMBXt motors allowed robust, high-resolution single-molecule tracking analyses. The hMBX motors undergo processive motility towards MT plus-ends over long distances with infrequent pauses (Figure 2E-F), while pauses are more frequent for the hMBXt motors containing the tail, similar to our observation with the kMBXt motors (Figure 2G-H). The hMBXt motors exhibited multiple intensities with bright and dim spots suggesting multi-motor clustering similar to the properties of kMBXt motors. Thus, for both *Drosophila* and human kinesin-5 mini-tetramer motors, the presence of the kinesin-5 C-terminal tail results in slower motors that undergo pauses, form clusters, and are more processive (Figure 2I, J).

### The kinesin-5 tail domain promotes static pausing

To examine kinesin-5 mini-tetramer pausing behavior at the single molecule level, we analyzed the hMBX motility tracks by sub-pixel localization and linked these into trajectories (Arcizet et al., 2008; Tinevez et al., 2017)(Hafner et al., 2016; Zajac et al., 2013). The mean-squared displacements (MSDs) were compared as a function of time on a log-log plot, where a slope of α=1 indicates a purely diffusive process, α<1 in the case of confined diffusion, and α=2 in the case of processive transport (Arcizet et al., 2008); (Hafner et al., 2016; Ruthardt et al., 2011; Zajac et al., 2013). We repeated this fitting process in a sliding window along the trajectory to calculate a local slope or α-value for each point in the trajectory (Figure 3A, Figure 3 figure supplement 1), and then used change-point analysis to identify segments of processive and paused motility (Beausang et al., 2011). The α-value fluctuated between ~0 and ~2, suggesting hMBX motors are either undergoing processive motility or remain tightly bound to single sites along MTs, without diffusing. We observe a bimodal α-value distribution with peaks at 0.627+/0.041 and 1.760 +/- 0.041 (Figure 3C). Our analysis suggests that hMBX motility changed significantly within a given trajectory between either paused or processive motile segments. The change point analysis likely underestimates of the number of short pauses. To account for pausing more effectively, we also fit the data for pauses ≥5 frames to a single exponential distribution, yielding a mean pause time of ~13.5 s (Figure 3 figure supplement 1). The pause time distribution is described well by a single exponential. Quantifying the number of pauses per trajectory revealed that ~41%, ~27%, and ~12% of long trajectories contained zero, one, or two pauses, respectively (Figure 3 figure supplement 1). To allow for more averaging and minimize the effect of statistical fluctuations, we next calculated the MSD for all of the hMBX motility segments identified as paused or processive. As the localization error is most prominent at short timescales (Michalet, 2011) and is expected to be of a magnitude similar to the displacement of the motor due to directed motion in our experimental conditions, we focused our analysis on timescales of ~3-7 seconds (Duselder et al., 2015), also avoiding errors due to limited averaging at longer timescales (Michalet, 2011). The hMBX motor MSD curves exhibit two unique α-values of 0.627 and 1.760 indicating either paused and processive motility trajectories, respectively (Figure 3C).

**Figure 3.**
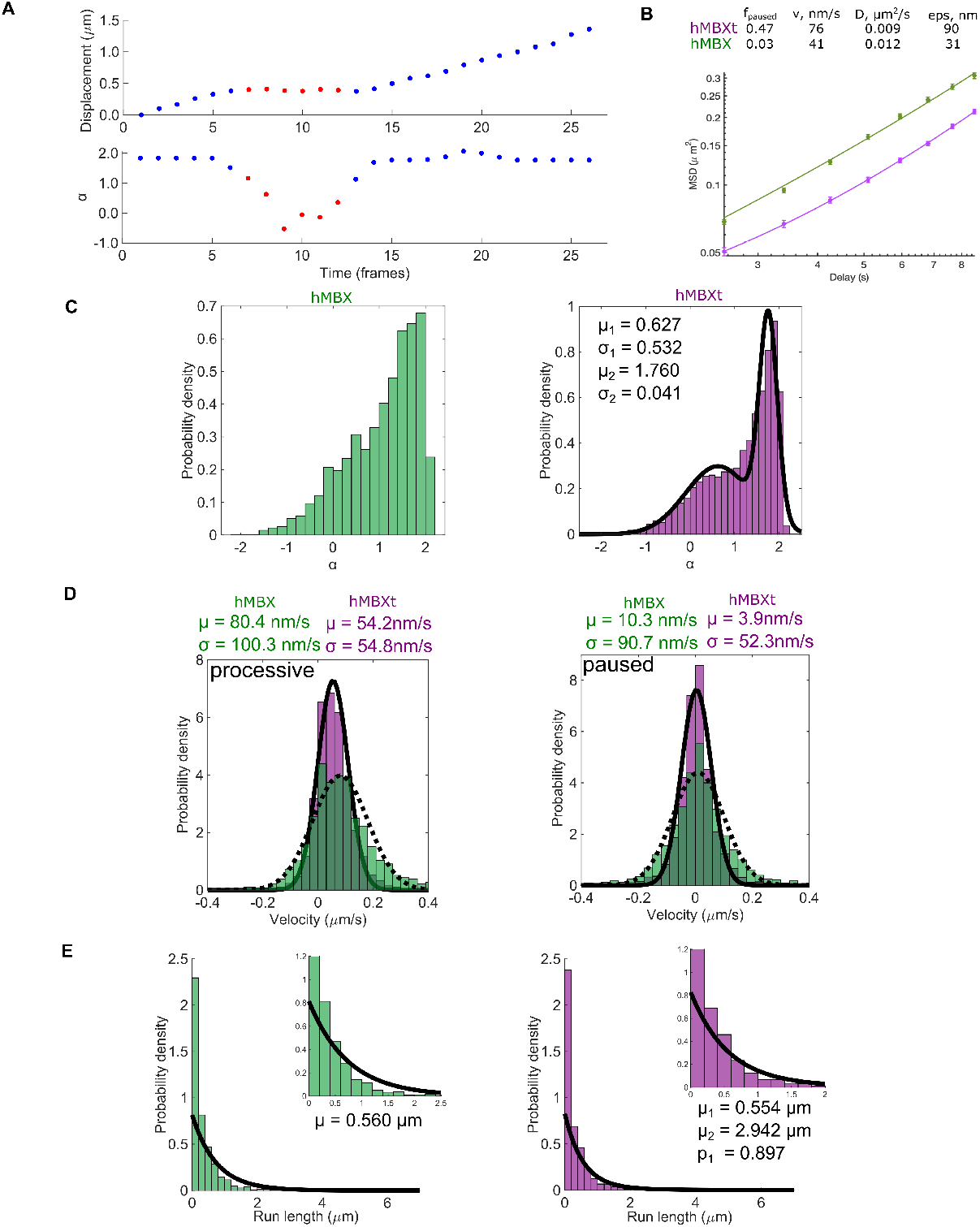
The Kinesin-5 tail domain induces pauses, increases run lengths and decreases motility velocities of the mini-tetramer kinesin-5. A) Pauses (red) and periods of processive motility for hMBX motor (blue) were identified in trajectories based on the slope of the mean-squared displacement (α) calculated within a sliding window along the trajectory. α fluctuates between ~2 and ~0 indicating processive and stationary motility (bottom). B) The MSD for the entire trajectories of hMBX motors were fit to an expression that describes periods of stationary pauses and motile events consisting of processive and diffusive movements, MSD=(ϕvt)^2+ϕ2Dt+2e^2 where ϕ is the fraction of time in a motile state. HMBX motors pause more frequently. C) Plotting all local α-values reveals primarily processive motility (green), while hMBXt exhibits a bimodal distribution with peaks consistent with static binding (μ1) and processive motility (μ2) (purple). D) Histograms showing the frame-to-frame velocity of hMBXt motors (purple; solid line) and hMBX motors (green; dotted line). hMBX motors move faster than hMBXt motors during the processive sections of trajectories. Velocities are similar during paused sections. (E) The total run length for hMBX motors was fit with a single exponential distribution with mean μ. All trajectories >3 frames were included (n = 2056 trajectories). Unless otherwise specified, n = 111 trajectories from 3 independent experiments for hMBX constructs and n = 143 trajectories from 9 independent experiments for hMBXt constructs.

To determine the effect of the human Eg5 tail on motility properties of mini-tetramers, we compared hMBXt to hMBX motility independent of the change point analysis, we also calculated the MSD for all of the hMBXt motility events (paused and processive segments) and fit it to the function < *r*(τ)^2^ >= (ϕ*ν*τ)^2^ + ϕ2*D*τ + 2ϵ^2^ where ϕ is the fraction of time in the processive state (Chugh et al., 2018). From this global analysis, the fraction of time paused (1 - ϕ) is 0.47 for hMBXt compared to 0.03 for hMBX motors (Figure 3B). These results indicate that pauses are due to a statically bound state of the hMBX motor. For processive sections, the hMBX motors α-value near 2 indicates that the motor’s motility is dominated by active transport. Thus, our analysis suggests that hMBX motor paused states represent a single state that is tightly bound to MTs without diffusion. The hMBX motor motility velocity during processive motility segments displayed a single distribution (Figure 3D) (54.2 ± 1.4 nm/s). The velocity distribution for the paused fraction peaked near 0 nm/s (μ = 3.9 ± 1.8nm/s and σ = 52.3 ± 1.3nm/s) suggesting clearly static motors that are strongly attached to MTs (Figure 3D).

Next, we analyzed the duration of hMBX processive motility events between paused states. We determined the distance travelled by a motor without pausing as the total run length. Since not all motile hMBX motors exhibited pauses, we observed two sub-populations: motors in which there was a pause, and motors in which the motility was uninterrupted. Correspondingly, distributions for total run lengths for hMBX motors were fit to a two-term exponential distribution, with run lengths of 554 ± 60nm and 2942 ± 1106nm, with 90 ± 5% of spots in the shorter run length population (Figure 3E). Plotting the inter-pause run lengths revealed a single exponential distribution with a mean run length of 1263 ± 167nm (Figure 3E, Figure 3 figure supplement 1). Thus, the shorter of the two estimates for total run lengths likely corresponds to the inter-pause run length or that of motors, which do not pause. This suggests that hMBXt motors may string together multiple shorter runs with pauses, allowing for a longer total run length. Both human and *Drosophila* mini-tetramers exhibit very similar pausing behavior (Figure 2C-D, G-H).

To confirm the static nature of the hMBX pauses, we performed motility assays with the slowly hydrolysable ATP analog, AMPPNP, which is expected to trap motors in a strongly bound state (Chen et al., 2016). As expected, hMBX exhibited only static binding in the presence of AMPPNP (Figure 3 figure supplement 1) with normally distributed α-value with a maximum near 0.103 and a standard deviation 0.316, indicative of static binding (Figure 3 figure supplement 1). AMPPNP trapped hMBX motors show static binding similar to pauses observed in the presence of ATP. Thus, we propose that hMBX and kMBX motors switch between processive and static bound states, while moving along MTs toward plus-ends. We propose these paused states represent the nucleotide-free state rather than the ATP bound state, in both of which the motor domain has high affinity for MT lattice sites (Cross and McAinsh, 2014).

### The kinesin-5 tail domain promotes motor clustering

Both kMBXt and hMBXt mini-tetramers formed multi-motor clusters along MTs during pauses (Figure 4A, B). Thus, we next analyzed the properties of these multi-motor clusters and how they form on MTs. First, we studied the role of electrostatic interactions in motor clustering and the impact of clustering on the motile properties of the motors. We compared kMBXt motor behavior across five ionic strength conditions (25, 50, 100,125,150 mM KCl) with the properties for kMBX motors in two ionic conditions (50 and 100 mM KCl) (Figure 4C). For each condition, we normalized the intensity of spots corresponding to motile motors along MTs visible in each field of view and analyzed the corresponding intensity distribution to determine the average intensity of individual mini-tetramer (Pandey et al., 2021)(Figure 4 figure supplement 1). The analysis revealed a major peak of spot intensity likely representing individual kMBXt mini-tetramers, but higher intensity spots were also observed, representing larger multi-motor clusters. We also determined the rate of kMBXt clustering formed during motility by measuring and quantifying the events in which motor intensities merged to form brighter spots, and quantified these events in the pools of motile motors in the different ionic strength conditions (Figure 4C). At 25-50 mM KCl, the majority of kMBXt motors formed clusters with either 2 (1+1) or 3(1+2), denoting the mini-tetramer ratio of motile motor to static motor (motile + static) components(Figure 4, figure supplement 2). However, at 50-150 mM KCl, a wider range of larger clusters was observed, representing 5-10 motors per cluster (1+4 to 2+10). We then calculated the total frequency for kMBXt cluster merging per unit MT length at 25-150 mM KCl, revealing the highest frequency peaks around 100 mM KCl, but this frequency is substantially lower at 25 mM or 150 mM KCl. In contrast few clustering events were observed for the KMBX motor in the absence of the tail domain at 50 and 100 mM KCl (Figure 4C). Thus, the kinesin-5 tail is responsible for kMBXt multi-motor clustering. At 25-50 mM KCl, clustering is likely low because the lower ionic strengths promote an intra-molecular tail-motor domain interaction due to their proximity. In contrast, at intermediate ionic strengths (100 mM KCl), the tail promotes interactions between multiple mini-tetramers and form clusters. At high ionic strengths (150 mM KCl), neither interaction is favored, again limiting the occurrence of interactions between mini-tetramers and thus diminishing the formation of clusters.

**Figure 4:**
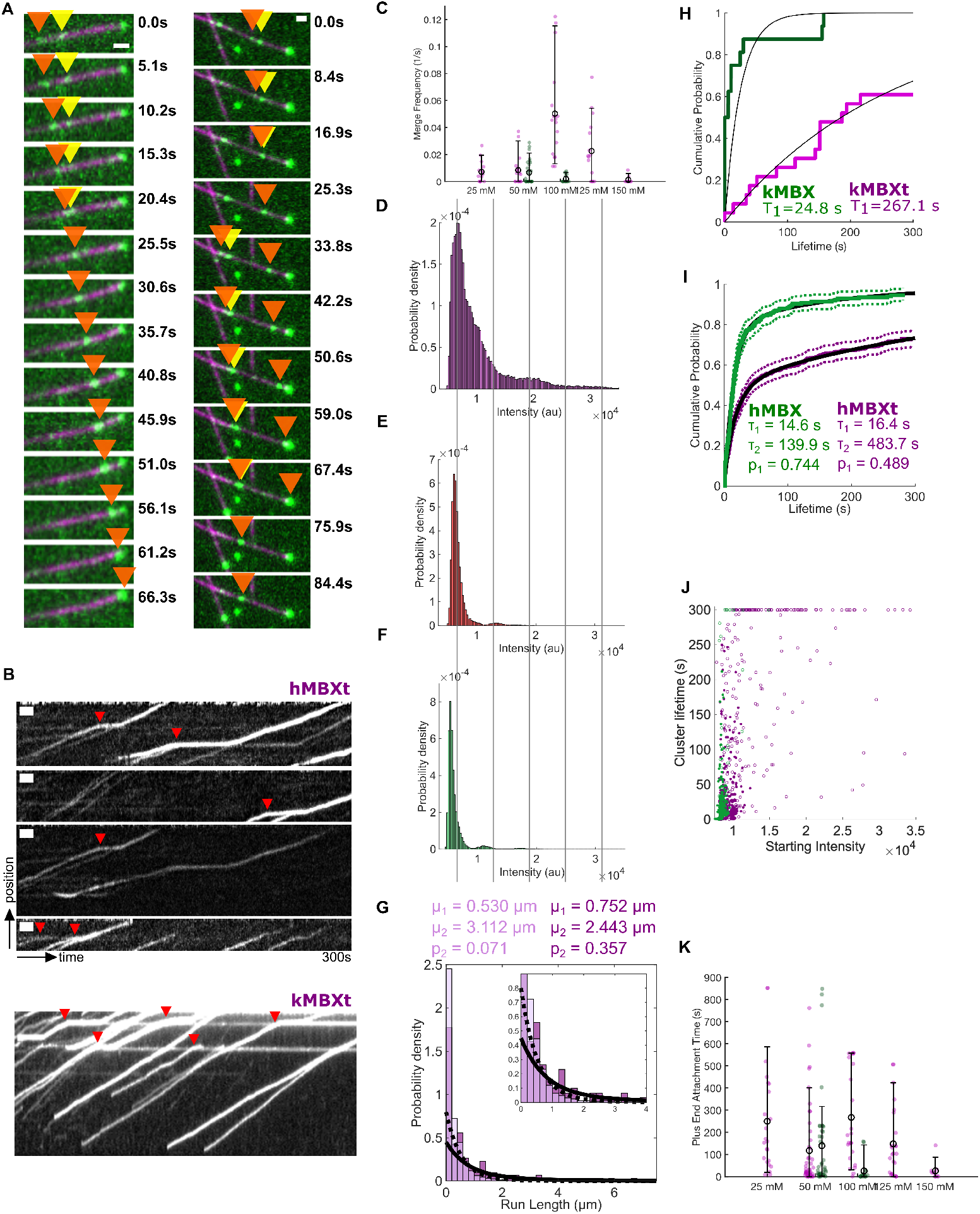
Kinesin-5 mini-tetramers with the tail domain, but not those without the tail, form multi-motor clusters while undergoing motility along a MT. A) Mini-tetramers with the tail form clusters while moving along the MT. Two such examples are shown. Often, one of the motors involved is paused (yellow arrow), a processively moving motor joins it (orange arrow), and the two continue moving together (orange) arrow). 6 (left) or 10 (right) frames between time points. Scale bars 1μm wide. B) Kymographs showing the formation of clusters. Note that many of these cluster’s form while one of the motors involved is paused (red triangles). Scale bars 10s wide and 1μm tall. C) The cumulative frequencies and their average for kMBX (green) and kMBXt (pink) motors to merge into clusters in relationship to the salt concentration (mM) conditions. The extended data for kMBX and kMBXt motor clustering are shown in figure supplement. D, E, F) Intensity distribution for constructs (D) with the tail domain in the presence of ATP, (E) with the tail domain in the presence of AMP-PNP, and (F) without the tail domain in the presence of ATP, suggesting that the mini-tetramers require the tail domain and motility in order to form clusters. Clusters likely correspond to 2-3 motors, and the blurring of the intensity distribution for motors in the presence of ATP is likely due to the quenching and unquenching of the TMR sensors used as labels during motor stepping. Lines are ~6500au apart. For D, n = 1406 trajectories from 9 independent experiments. For E, n = 1298 trajectories from 2 independent experiments. For F, n = 2056 trajectories from 3 independent experiments. G) Total run length distributions for single motors (light purple; dotted line) and clusters (dark purple; solid line), both with the tail. Distributions were fit with a double exponential distribution, and both groups have a short (μ1) and long (μ2) run length population. However, the fraction of motors in the long run length population (p2) is larger for clusters. H, I) Motors often reach the plus-ends of stabilized MTs and remain bound there for some time. Lifetimes were fit to a Weibull distribution. The hMBXt motors were best fit by a double Weibull distribution, suggesting the presence of populations with a short and long lifetime. The short lifetime (τ1) was similar for both constructs, whereas the long lifetime (τ2) was longer for hMBXt than hMBX motors. The fraction of the population with a short lifetime (p1) was also higher for hMBX motors, agreeing with the observation that hMBX motors do not form clusters along the length of the MT. J) Distribution of lifetimes of motors/clusters at the ends of MTs as a function of their starting intensity (average intensity of the first 5 frames they are detected). hMBX motors (green) generally arrive as single motors, whereas hMBXt motors (purple) often arrive as clusters. Open circles denote censored lifetimes (present in first or last frame of movie, indicating they are at least as long or longer than the point plotted). K) The cumulative MT plus-end association times for kMBX and kMBXt motors and their average values in relationship to the salt concentration (mM). The extended data for kMBX and kMBXt motor clustering are shown in Figure 4 figure supplement 2-3.

Next, we studied the relationship between kMBXt motor cluster size and average velocity, run length and run time. At 25 and 50 mM KCl, run length and run time are shorter and correlated with fewer motors per cluster. In contrast, at 100 and 125 mM KCl we observe an 80-100% increase in run length and run time and these features correlate with the larger motor-clusters. However, we found no direct correlation between cluster size and an increase in run length or run time within a single pool of motile kMBXt motors. Thus, our data demonstrate a critical role for the kinesin-5 tail mediated clustering in promoting slower motility and tight association with MT-plus ends. At lower ionic strength kMBXt motors behave more as individual tetrameric motors with shorter run lengths. At 100 mM KCl, kMBXt motor clustering becomes dramatically enhanced leading to motors that undergo motility more slowly with longer run lengths, run times.

The hMBXt motile motors similarly formed clusters with paused hMBXt motor along MTs (Figure 4A-B, Figure 4 figure supplement 1-3). To estimate more precisely the number of hMBXt motors in these clusters, we examined their spot intensities using multi-modal distribution analysis, revealing with a broad peak at ~6500au and peaks at 2 and 3 times this value (~13000au, ~19500au) (Figure 3D). The major peak likely represents individual hMBXt mini-tetramers, with the other peaks representing approximately 2 or 3 mini-tetramers. We tested the number of hMBXt motors in these clusters in the presence of AMPPNP, which led to non-motile attachment events. The spot intensity distribution for hMBXt in the presence of AMPPNP was predominantly unimodal (major peak at ~6100au and minor peak at ~13000au), suggesting that motility is required for hMBXt clusters to form along MTs (Figure 4E). In the presence of AMPPNP (i.e. without motor stepping), the major and minor peaks were clearly segregated; however, in the presence of ATP, these peaks were far less distinct (Figure 4D). This is likely because of the quenching and unquenching of the fluorophore pairs occurring with each step of the motor (multiple times during each frame) (Toprak et al., 2009). The multi-modal spot intensity distribution of hMBX motors was similar to that of the hMBXt motors in the presence of AMPPNP, with a major peak at ~5300 au (Figure 4F). The small shift in intensity is likely due to a difference in TMR labeling ratios for the two constructs.

Next, we measured the run length of hMBXt motor clusters moving along MTs (Figure 4G). Quantifying the total run length of “clusters” (spots with an intensity ≥1.5-fold higher than the mean intensity for single hMBXt tetramers) revealed run lengths of 752 ± 385 nm and 2443 ± 1649 nm, similar to the run lengths of 530 ± 58 nm and 3112 ± 1468 nm for spots likely representing single hMBXt motors (Figure 4A). However, only 7 ± 4% of single hMBXt motors were in the longer lifetime population, whereas 36 ± 40% of hMBXt clusters were in this population. Additionally, many clusters reached the plus-ends of the MT on which they were moving, remaining bound at the MT plus-end. For very processive motors, the MT length has an effect on the measured run length distribution because many motors reach MT ends such as their runs are artificially truncated (McHugh et al., 2018; Thompson et al., 2013). Thus, we accounted for this censoring when fitting the run length distribution. Notably, satisfactory fits to either of the run length distributions, for either single motors or clusters could not be obtained using single exponential distributions, indicating that there is likely an inter-pause and total run length. Thus both single motors and clusters are able to string together runs with pauses.

We subsequently analyzed the accumulation of hMBXt and kMBXt motors at MT plus ends. The hMBXt and kMBXt motors are highly processive, with run lengths that depend on the lengths of MTs, since many motors reach MT ends (McHugh et al., 2018). We observe an accumulation of both hMBXt and kMBXt motors at MT plus-ends, indicative of the stable association of tail-containing motors with MTs (Figure 4A). We quantified the dwell times of hMBXt motors at MT plus ends by measuring the intensity in a region centered at the plus-end of each MT and defining the period of time for which the intensity was >3 standard deviations above the background signal as the cluster lifetime. The tail domain enables mini-tetramers to maintain association at MT plus ends, where hMBXt motors pause for ~10x as long as hMBX motors (Figure 4H, I). For hMBXt, we could not obtain a satisfactory fit to cluster lifetimes using a single Weibull distribution, prompting us to use a mixed Weibull distribution with dwell-times of 16.4 ± 4.9 seconds and 483.7 ± 153.2 seconds (Figure 4I, Figure 4 figure supplement 5). This suggests the presence of two populations of plus-end clusters with different dwell times, with 49 ± 15% of motors in the shorter lifetime population. Next, we analyzed cluster lifetimes as a function of the intensity of the first 5 frames of that end cluster, revealing that motors that arrived in a cluster that had “pre-formed” along the MT were likely to remain associated with the plus-end (Figure 4J). We used this measure of the starting spot intensity rather than the mean spot intensity to distinguish between motors arriving in “pre-formed” clusters and motors arriving individually and gradually forming a cluster at MT plus-end. The arrival of “pre-formed” clusters was supported by the observation that intensity traces in the region near MT plus-ends generally increased above baseline or decreased to baseline in one or a few frames, regardless of the plateau intensity (Figure 4J, Figure 4 figure supplement 6). Thus, the shorter lifetime likely represents the dwell time of single hMBXt motors at the ends of MTs, while the longer lifetime likely represents that of clusters.

The hMBX motors did not accumulate as robustly at MT plus-ends (Figure 4I). The dwell times of hMBX motors at the plus-ends of MTs were 14.6 ± 3.2 seconds and 139.9 ± 137.5 seconds, with 74 ± 18% of motors in the shorter lifetime population (Figure 4I, Figure 4 figure supplement 4). Thus, the shorter dwell times for hMBX and hMXBt clusters are similar, in agreement with the idea that this represents the lifetime of single motors (present for both constructs) at MT plus-ends. In contrast, the longer lifetimes are shorter for hMBX motors, and a smaller fraction of motors is likely to be in this longer dwell-time subpopulation, indicating that the inability of hMBX motors to cluster may prevent them from remaining bound at MT plus-ends for extended periods (Figure 4H, I, J). To determine the possibility of hMBXt clusters reaching MT plus-ends, we next compared the distribution of frame-to-frame velocities for single motors and clusters in the paused and processive states (Figure 4). Our analysis agrees with the visual observations, that hMBXt clusters move more slowly than single hMBXt motors: whereas the mean paused velocities of single motors (4.6 ± 3.1nm/s and σ = 64.8 ± 2.2nm/s) and clusters (3.6 ± 1.9nm/s and σ = 37.6 ± 1.3nm/s) were similar, the processive velocity of single motors (63.3 ± 2.3nm/s) was higher than that of clusters (43.2 ± 1.6nm/s). Additionally, the velocity of single motors was more variable than that of clusters. During the processive sections of trajectories, the standard deviation of single motor velocities was 65.3 ± 1.6nm/s, whereas that of clusters was 39.5 ± 1.2nm/s. This might contribute to the observation that the velocity of hMBXt motors (54.2 ± 1.0nm/s) is slower than that of hMBX motors (80.4 ± 4.4nm/s).

We also measured the average dwell time for ensembles of kMBX and kMBXt motors at MT plus ends in different ionic strength conditions. We observe that kMBX motors dwell at MT ends for 78 ± 19 s at 50 mM KCl and for 25 ± 18 s at 100 mM KCl, showing a three-fold decrease upon ionic strength increase (Figure 4K). In contrast kMBXt motors show consistently longer dwell time at MT plus-ends, which is two-three folds higher than kMBX motors (125 to 250 s), and are not influenced by changes in ionic strength (25-150 mM KCl) (Figure 4K). The majority of kMBXt clustering events also correlate with the motors arriving at MT plus-ends particularly at 100 mM KCl condition. This is likely due to the enhanced multi-motor clustering coupled with the enhanced dwell time at MT plus-ends at 100 mM KCl. Our data suggests that the MT plus-end dwell time is likely related to the motors pausing at MT plus-ends, and that the tail enhances this property by down-regulating the ATP hydrolysis activity of the motor domain.

### Short kinesin-5 mini-tetramers show unique MT crosslinking and MT sliding features

We next sought to determine whether kinesin-5 mini-tetramers are capable of crosslinking, aligning, and sliding pairs of MTs. To do this, we immobilized taxol-stabilized HiLyte 646 and biotin labeled MTs on a coverslip via a streptavidin-biotin linkage. We found that to achieve efficient MT bundling, particularly for the mini-tetramer motors, we required high velocity flow of MTs in flow cells such that they were well nearly aligned with the direction of flow. Next, motors were introduced into the sample chamber at a specified concentration and allowed to decorate the coverslip-attached MTs. Finally, Rhodamine-labeled MTs and ATP were introduced with rapid flow rates, and kinesin-5 mediated crosslinked MT bundles were allowed to form and motors to initiate relative MT sliding. We then imaged both the coverslip attached MT, paired MTs and kinesin-5 motors using the three different channels via TIRF microscopy at 2-5 sec frame rates.

We first set out to determine how efficiently four different kinesin-5 motor constructs can crosslink and pair MTs, by comparison: full-length Eg5 with tail (hFLt), full-length Eg5 without the tail (hFL), previously prepared as described (Bodrug et al., 2020), kinesin-5 mini-tetramer with tail (kMBXt), and kinesin-5 mini-tetramer without tail (kMBX). We then imaged multiple fields-of-view immediately after flowing in ATP and Rhodamine-MTs and measured how many MT bundles had formed relative to the population of surface-bound MTs (Figure 6a-b). We found that both hFLt and hFL motor constructs recruited free MTs from solution, forming MT pairs that underwent sliding at relatively high rates, with 63±5% and 54±16% in the hFLt and hFL motor conditions, respectively. This consistent with observations we made previously at similar concentrations (Bodrug et al., 2020). In contrast, the kMBXt motor formed MT bundles 3-fold less frequently (24±14%) compared to hFLT, while the kMBX almost never recruited a free MT from solution and rarely formed MT pairs, with only 2% of all surface MTs observed to bundle a second MT (Figure 5A-B). Together, these data suggest that the kinesin-5 mini-tetramer motors are less efficient than the full-length kinesin-5 motors at spontaneously forming antiparallel MT bundles and that the tail domain is critical for establishing the MT crosslinked geometries, especially in the mini-tetramer constructs.

**Figure 5:**
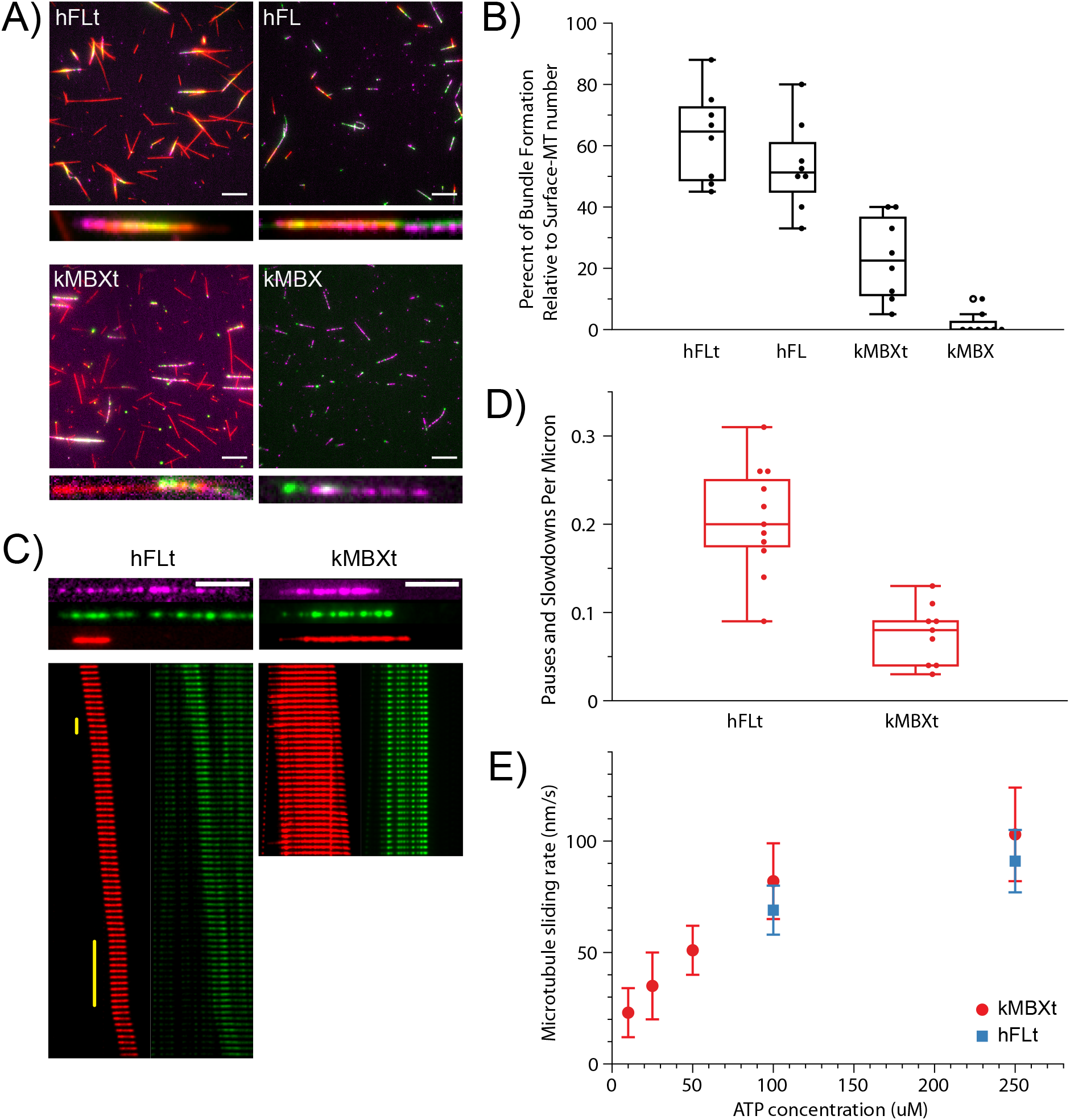
Kinesin-5 mini-tetramers show defects in crosslinking MTs but enhanced MT sliding once aligned, compared to native kinesin-5. A) Representative TIRF images of surface-immobilized MTs (magenta) crosslinked via kinesin-5 (green) constructs to free MTs (red). Four different kinesin-5 constructs were examined: hFLt, hFL, kMBXt, and kMBX. Scale bar = 10 microns. B) Percentage of surface-immobilized MTs that engaged in kinesin-5 mediated crosslinking with free MTs for each kinesin-5 construct. N=8 fields of view analyzed per condition. C) Sample MT pairs (top) and kymographs (bottom) depicting MT sliding driven by either hFLt (left) or kMBXt (right). Free MT (red) and kinesin-5-GFP or nNG (green) are shown. Pauses are identified by vertical yellow bar. Scale bar = 4 microns; frame rate for hFLT = 10 seconds per frame; for kMBXt = 5 seconds per frame. D) Number of pauses or velocity reduction events observed per micron for hFLt and kMBXt driven MT sliding. N = 11 events for hFLt, N = 9 events for kMBXt. E) Average MT sliding rate calculated for bundles at different ATP concentrations. N = 6 events for each condition. Error bars are S.D. Values are reported in Table 3.

We next sought to determine how kinesin-5 mini-tetramers compared to full-length kinesin-5 in MT crosslinking, pairing or alignment and then MT sliding. We therefore monitored the positions of the mobile MT and the full length or mini-tetramer kinesin-5 motors during these sliding events across a range of conditions. As we were unable to form bundles using kMBX motor in our assays, we focused on comparing the mechanics of the full-length (hFLt) motor and mini-tetramer motor (kMBXt) both of which contain kinesin-5 tail domains. Both motors were able to slide MTs apart efficiently (Figure 5C). Interestingly, kMBXt motor appeared less mobile and more clustered relative to the hFLt motor along the MTs throughout MT sliding events (Figure 6c). We also observed that the mobile free-MT occasionally paused or exhibited reduced velocity for brief stretches when undergoing sliding by the full-length kinesin-5 (hFLt). In contrast, MT sliding generated by kMBXt motor tended to exhibit more consistently continuous MT sliding motility with faster velocities throughout (Figure 5C). To examine this difference, we determined the frequency of observed pauses or velocity decreases across many MT sliding examples and found that MT bundles in the full length kinesin-5 (hFLt) condition exhibited pauses 0.21+/-0.06 times per micron (μm) of distance travelled, while kinesin-5 mini-tetramer (kMBXt) paused at a lower rate of 0.08+/-0.04 times per micron (μm) which is three-folds lower than the pausing exhibited by full length (hFLt) motor MT sliding (Figure 5D). Finally, we calculated the MT sliding velocity across a range of ATP concentrations for both constructs (Figure 5E). We found that the MT sliding velocity was approximately twice the speed of single motor stepping motor velocity, consistent with the previously reported finding that both pairs of motor domains at each of the bipolar end of kinesin-5 motors are stepping along their respective MT in the opposite direction, leading to a two-fold MT sliding motility compared to the motility generated along single MTs (Kapitein et al., 2005). Importantly, we observed that the average velocity of kinesin-5 mini-tetramer (kMBXt)-driven MT sliding at 100 and 250 μM ATP was about 10% higher than that of full-length kinesin (hFLt)(Table 3). Together, these results indicate that, while the kinesin-5 minitetramers are not as efficient at initially crosslinking aligning two MTs as native kinesin-5, they are capable at sliding MTs, doing so within clusters, with less three-fold less pausing and 10% higher velocities compared to those generated by full-length kinesin-5.

**Figure 6:**
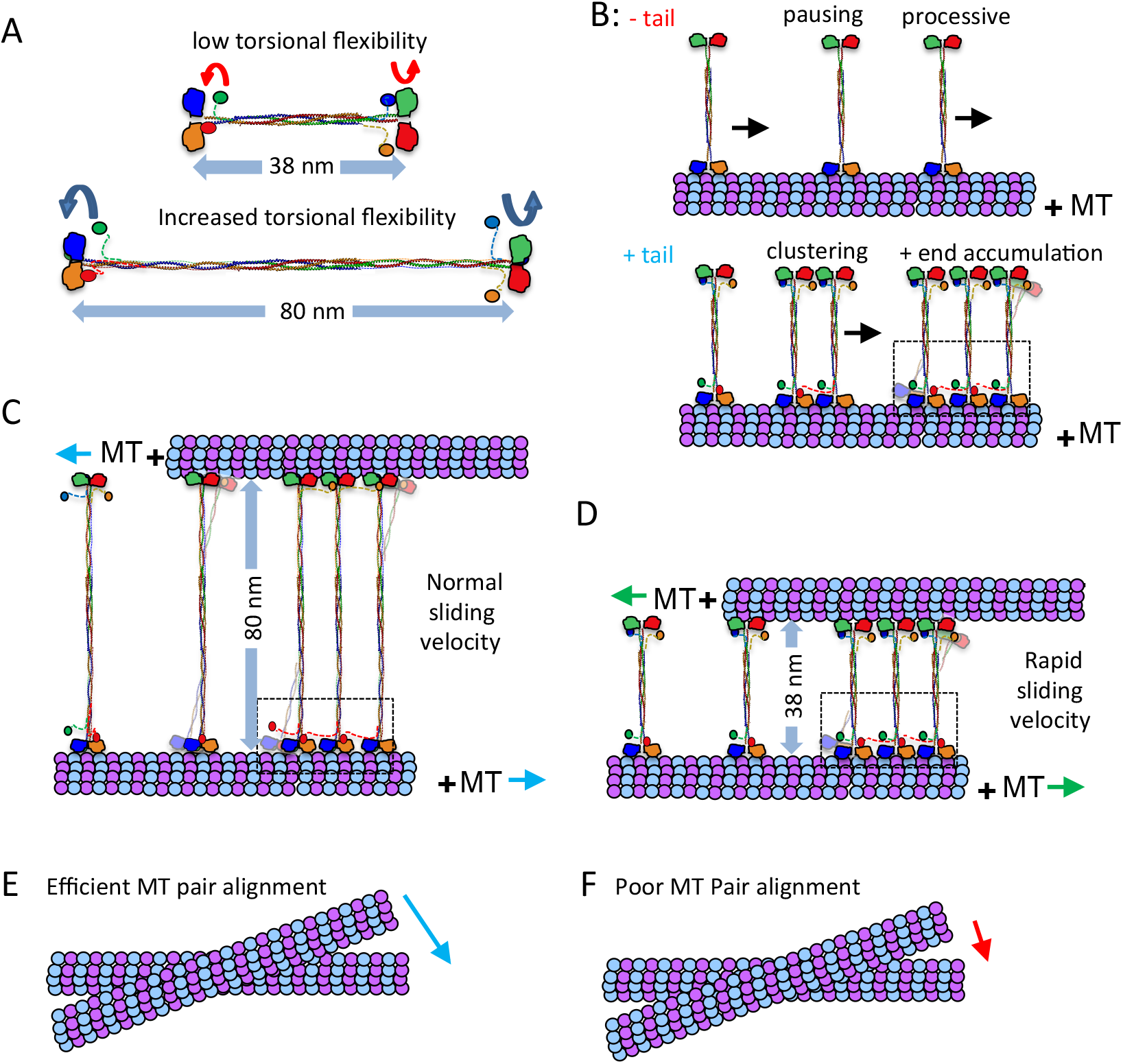
Kinesin-5 tail to motor regulation and minifilament length modulate motor clustering and features of MT-sliding motility. A) The Kinesin-5 mini-tetramers are 38 nm in length leading to a decrease in torsional flexibility compared to native kinesin-5, which are 80 nm in length and have higher torsional flexibility. B) Top panel, kinesin-5 mini-tetramers without the tail domain show processive motility punctuated by pauses and little residence at MT-plus ends. Bottom panel, kinesin-5 minitetramers with the tail domain show increased pausing, coupled with motor clustering mediated by cross motor-tail interactions between mini-tetramers. C) MT sliding motility mediated by native kinesin-5 leads to 80-nm separation between paired MTs, which are slide apart with normal sliding velocity, which is punctuated by pauses and is lower than twice motility velocity of each motor end along each MT. D) MT sliding motility by the kinesin-5 mini-tetramers leads to 38-nm separation between the paired MTs and a more efficient MT sliding motility that approaches closely to twice the motility velocity of each motor end. E) The native kinesin-5 motor MT pair alignment is efficient due to torsional flexibility of the minifilament F) Kinesin-5 mini-tetramer MT pair alignment is poor due to decreased torsional flexibility of its shortened minifilament.

**Table 3:** MT sliding velocities for Native and mini-tetramer kinesin-5 motors

| ATP ($\mu\text{M}$) | hFLt | kMBXt |
| --- | --- | --- |
| 10 | N/A | 23 +/- 11 nm/s (n=9) |
| 25 | N/A | 35 +/- 15 nm/s (n=6) |
| 50 | N/A | 51 +/- 12 nm/s (n=11) |
| 100 | 69 +/- 11 nm/s (n=13) | 82 +/- 17 nm/s (n=15) |
| 250 | 91 +/- 14 nm/s (n=17) | 103 +/-21 nm/s (n=14) |

## Discussion

Human cells encode 45 kinesin motors, each of which is adapted to perform a unique set of cellular functions. This functional diversity is demonstrated beautifully in the mitotic spindle, where kinesins perform diverse tasks ranging from regulating dynamics to crosslinking and sliding MTs. To fulfill their functional niches in mitosis, kinesins have evolved biophysical and biochemical differences that render them uniquely well-suited for these roles. Through comparing the motility and structure of full-length and minimal tetrameric kinesin-5 motors, our studies reveal the role of the C-terminal tail and bipolar tetrameric minifilament domains in governing kinesin-5 motor clustering and MT sliding motility. The structural and biochemical features of these domains are highly conserved across all the kinesin-5 motors. Our structure of the extended kinesin-5 BASS domain reveals that it possesses the unique capacity to form a force-bearing junction for two pairs of motors positioned at opposite ends of its bipolar structure. We designed and studied Human and *Drosophila* kinesin-5 mini-tetramer motors based on the BASS-XL x-ray structure (Figure 1), which recapitulate critical aspects but reveal unique differences compared to full-length kinesin-5 motor motility (Bodrug et al., 2020). These structural adaptations are critical for kinesin-5 function in bipolar mitotic spindle assembly, organization, and elongation.

### The kinesin-5 tail domain binding regulates the motor domain mechanochemical cycle and promotes slower motility and increased processivity

The kinesin-5 tail domains down-regulate MT activated motor domain ATP hydrolysis by stabilizing the MT bound nucleotide free state. This interaction leads to slower stepping during processive motility, longer run lengths, and frequent pauses (Figure 2–3). As with full length kinesin-5 (Bodrug et al., 2020), the presence of the tail domains in both *Drosophila* and human mini-tetramers leads to slow motility with multiple pauses in which motors are statically bound to MTs (Fig. 2). The pauses represent strongly-bound states with extended lifetimes in which the leading motor domain is in the no nucleotide state, while and the trailing motor domain is either in the ADP-P_i_ or ADP state, similar to the so-called ATP gate or stepping gate (Andreasson et al., 2012; Bodrug et al., 2020; Cross and McAinsh, 2014). This is similar to motor motility in the presence of a mixture of ATP and AMP-PNP, in which comparable switches in motility are observed (Subramanian and Gelles, 2007; Vugmeyster et al., 1998). We suggest that the kinesin-5 tail domain docking onto the motor domain serves as an externally imposed gate to organize multiple kinesin-5 motors in clustered assemblies. The kinesin-5 tail domain enhances the pausing and promotes motor assembly into clusters via encounters between motile and paused motors along MTs. The tail to motor trans-interactions between the tails of one kinesin-5 tetramer and the motor domains of other tetramers. Interestingly, motor clustering has also been described for the budding yeast, *Saccharomyces cerevisiae*, ortholog Cin8 and has been proposed to mediate its minus-end to plus-end MT motility directionality reversal (Pandey et al., 2021a; Pandey et al., 2021b; Shapira et al., 2017; Singh et al., 2018).

### The kinesin-5 tail domain drives the formation of multi-motor clusters with different motile properties than single motors

Our studies of *Drosophila* and human kinesin-5 mini-tetramers reveal that tail to motor domain interaction is highly conserved, in which the tail domains induce pausing and drive kinesin-5 motors to assemble into multi-motor clusters. By increasing the pause frequency, the tail domain increases the frequency of encounters between motile and paused motors, allowing for the formation of multi-motor clusters mediated by trans-interactions between the tails of one tetramer and the motor domains of other tetramers (Bodrug et al., 2020). The tail to motor interaction leads to a decrease in overall motility velocities. Within clusters, multiple motors move together as assemblies (Figure 4). The tail may stochastically dissociate from the front motor domain, allowing the motor domains to continue stepping. The increase in the number of motors in these clusters likely leads to longer run lengths likely due to the increase in the numbers of stepping motors domains. Indeed, we observed an increased total run lengths in kMBXt and hMBXt motors compared to kMBX and hMBX motors, supporting the effect of the tail on increasing processive run lengths. The lifetime of motors at MT plus-ends is related to the number of motors in the clusters that arrive. The kMBX or hMBX motors generally dissociate from the MT plus-end more quickly than clusters of kMBXt or hMBXt motors (Figure 4). Additionally, kMBXt or hMBXt motor clusters were found to move at slower and less variable speeds than single motors (Figure 4). Clusters and single motors may be fulfilling different roles during spindle assembly. Clusters of motors may be present between parallel MTs, such as near the spindle poles, where, because of their increased association time with MTs, they could assist in MT capture early in spindle assembly. Kinesin-5 clusters may be selectively retained in this region because they move slower than single motors (Fig. 4). In contrast, faster moving single motors may “escape” and travel towards MT plus-ends, localizing them in the region of antiparallel MT overlap in the spindle midzone. Here, they could facilitate spindle pole separation and/or regulate the rate at which this occurs

### The kinesin-5 tail-motor interaction is likely tuned by mitotic phosphorylation of the conserved BimC box in the tail domain

The kinesin-5 tail domain contains a conserved BimC box or CDK1 phosphorylation site (Threonine 926 in human kinesin-5) and the more distal regulatory regions of the tail domain (i.e. the KEN box, D box, and Nek6 phosphorylation site (Blangy et al., 1995)(Bertran et al., 2011; Drosopoulos et al., 2014; Rapley et al., 2008). In particular, the phosphorylation of the kinesin-5 tail by Cdk1 kinase has been shown to increase the affinity of the motor for MTs *in vitro* (Cahu et al., 2008). Based on our observations, we hypothesize that such phosphorylation may enhance the tail-motor interaction by adding a negative charge to the tail, potentially increasing its affinity for the motor domain, which is positively charged near the ATP binding site. The CDK1 phosphorylation of the tail domain could act to increase the affinity of the motor for MTs by promoting tail docking, hereby increasing the time that the motor spends in its strongly-bound state. The enhancement of the tail regulation may lead to changes in the MT sliding mechanism or enhancing the brake function for kinesin-5 during anaphase. The impact of phosphorylation of the kinesin-5 tail warrants further study.

### The tail to motor interaction supports kinesin-5’s function as a brake during mitosis

Kinesin-5 motors have been suggested to act as a brake to slow the rate of MT sliding by other motors, both during anaphase (Rozelle et al., 2011; Saunders et al., 2007) and in non-mitotic cells (Falnikar et al., 2011; Lin et al., 2011; Myers and Baas, 2007; Nadar et al., 2008; Nadar et al., 2012). Moreover, the ability of kinesin-5 motors to act as a brake between parallel MT or quickly sliding antiparallel MTs has been demonstrated *in vitro* (Shimamoto et al., 2015). The tail docking onto the motor domain to induce pausing and motor clustering may contribute to the capacity of kinesin-5’s brake-like behavior by increasing the time these motors spend in their strongly bound state on the MT lattice. This would allow the motor to prevent rapid sliding of two MTs in a parallel orientation. The tail’s regulation is critical for optimal force transmission through kinesin-5 minifilaments to promote efficient crosslinking and sliding. We expect that a motor is less likely to detach when subject to loads while in tail-induced clustered and paused states than while moving processively as individual tetramers. The paused state of clustered motors would be particularly strongly attached to MTs, allowing them to withstand substantial loads when acting as a brake.

### Clusters of Kinesin-5 remain stably associated with the plus-ends of stabilized MTs

Kinesin-5 clusters often reached MT plus ends and they remained bound for extended periods of time (Figure 4). The tail to motor interaction strongly enhances the kinesin-5 the MT plus-end association. The accumulation of kinesin-5 at MT plus-ends has been described previously (Chen & Hancock, 2015; Kapitein et al., 2005). We interpret this MT plus-end accumulation to be due to the extended lifetime of kinesin-5 strongly-bound motor states, compounded by tail to motor interaction enhancing the binding affinities of the motors and enhancing their clustering (Furuta et al., 2013; Vershinin et al., 2007). Indeed, we observed that the motors that arrived at MT ends in clusters remained bound for longer periods of time (Figure 6c). In contrast to previous reports employing dimeric kinesin-5 constructs, it has proposed that this MT plus-end accumulation reveals a role for kinesin-5 motor domain in modulating MT dynamics (Chen et al., 2019; Chen and Hancock, 2015). However, we find that kinesin-5 mini-tetramers lacking the tail domain have short life times at MT plus ends, likely due to tail to motor interaction stabilizing the high affinity motor states and inducing multi-motor clusters. Furthermore kinesin-5 cluster also move with a slow velocity of ~43nm/s, which may be too slow to allow them to accumulate at the ends of dynamic MTs within the spindle.

### The length of the Kinesin-5 central minifilament directly regulates force transmission during MT sliding motility

Our studies help to elucidate the crucial role of the 60-80 nm Kinesin-5 minifilament in the kinesin-5 MT sliding mechanism. We compared kinesin-5 mini-tetramers with full-length kinesin-5 motors to determine the relationship between the length of the kinesin-5 minifilament to the efficiency of MT pair formation, and its impact on MT sliding motility activities. The kinesin-5 mini-tetramers are half the length of native kinesin-5 motors. The short minifilaments in kinesin-5 mini-tetramers are stiffer due to the shorter dimeric coiled-coils on either side of the BASS domain. This leads to low torsional flexibility and therefore enhance the force coupling between the bipolar ends of kinesin-5 as they engage the two MTs they crosslink. However, the decreased flexibility either end of the kinesin-5 mini-tetramers appears to impede efficient MT alignment into bundles. In particular, this decreased flexibility leads to defects in the initial crosslinking of MT pairs prior to MT sliding. Once crosslinked and aligned, however, the shorter minifilament leads to a higher maximal MT sliding velocity compared to native kinesin-5 due to the increased stiffness of kinesin-5 minitetramers.

Our studies also show the relationship between the kinesin-5 minifilament length and the motor to tail regulation in modulating MT sliding motility. The decreased kinesin-5 minifilament torsional flexibility in the mini-tetramer motors leads to dependence on the tail-motor regulation during MT crosslinking, alignment and sliding in contrast to full-length kinesin-5, where the tail was not strictly required *in vitro*. The kinesin-5 mini-tetramers show a very poor capacity to assemble paired sliding MTs in contrast to the full-length kinesin-5 motors such as the hFL (Δtail kinesin-5), which assembles MT sliding pairs, but with decreased efficiency relative to the hFLt (full length kinesin-5)(Bodrug et al., 2020). These data also show that the tail-motor regulation enhances force transmission properties of kinesin-5 engaged motor ends, and that the motility resulting forces are transmitted between kinesin-5 bipolar ends via their central tetrameric minifilaments.

Taken together, our studies identify how the unique structural adaptations of the kinesin-5 tail and tetramerization domains enable its role in organizing the mitotic spindle. The tail domain promotes motor pausing and clustering. Single motors and clusters behave differently, suggesting that they may fulfill different roles in spindle assembly. The length of the minifilament contributes flexibility to enable kinesin-5 to initiate MT crosslinking efficiently.

## Acknowledgement

The authors thank Dr Jonathan Scholey (Molecular Cellular Biology, UC-Davis) for the inspiration on the project. J.A.B thanks Dr. Richard Mckenney (Molecular Cellular biology, UC-Davis) for the critical reading of this manuscript. BASS-XL diffraction data were collected at the Stanford Radiation Laboratory (SSRL) using 11-1 beamline. We thank Peter Dunten, Ana Gonzalez, and Tzanko Doukov (SSRL) for help with data collection. This research was supported in part by the Israel Science Foundation grant (ISF-386/18) awarded to L.G.; the National Science Foundation (NSF-1615991) and United States - Israel Binational Science Foundation grant (BSF-2015851), awarded to L.G. and J.A.B., respectively; A United States - Israel Binational Science Foundation grant (BSF-2019008), awarded to L.G. and S.R.; and the National Institutes of Health (NIH-GM11283), awarded to J.A.B. S.F is supported by the Rensselaer Polytechnic Institute School of Science Startup funds. A.G.H is supported by the Canadian Institutes of Health Research (CIHR) grant PJT-159490 and the Natural Sciences and Engineering Research Council of Canada (NSERC) grant RGPIN-2020-04608. Crystallographic data and Structure coordinates are deposited at the RCSB (PDB ID: 7S5U).

## Materials and Methods

### Protein Production, X-ray crystallography and Model building

The KLP61F minifilament extended region (residues 597-833) was expressed and purified in BL21 *E. coli* as described in Scholey et al 2014. Bacterial pellets were lysed using a microfluidizer in (300 mM KCl, 50 mM HEPES, 1 mM MgCl_2_, 3 mM β-mercaptoethanol with protease inhibitors). The bacterial lysate was clarified by centrifugation at 18k rpm for 30 min at 4°C. Ni-NTA affinity was used to purify BASS-XL, and passage over HiTrap Q HP cation exchange in low salt (70 mM KCl, 50 mM HEPES, 1 mM MgCl_2_) was used to remove contaminants where BASS-XL eluted in the flow through. A second Ni-NTA affinity step was used in conjunction with 10K Amicon Filters to concentrate the BASS-XL. The concentrated BASS-XL tetramer was applied on a HiLoad 16/600 Superdex 200 gel filtration column using an AKTA Purifier (GE Healthcare). Crystallization conditions were screened using a Mosquito Robot (TTP Labtech) by mixing a 100 nL of protein with 100 nL precipitant conditions. Crystals were obtained and refined in 0.01 M FeCl3, 0.1M sodium citrate pH 5.6, 12% Jeffamine M-600 at 18°C and cryoprotected with 20% glycerol. Crystals were diffracted at the SSRL 11-1 beamline and showed highly anisotropic x-ray diffraction. Crystals adopt space group C2 with four molecules in the asymmetric unit. We used 4.4 Å as the high-resolution cut-off to avoid excessive loss of completeness. The diffraction data was truncated using boundaries determined via the Anisotropic server (Strong et al., 2006). The BASS-XL structure was determined using molecular replacement using the previously determined BASS model (Scholey et al 2014). Data from each monomer were combined using non-crystallographic symmetry and were averaged and refined using PHENIX with cycles of model building using coot program (Emsley et al., 2010; Liebschner et al., 2019). The individual positional coordinates and anisotropic B-factor were refined with automatic weight optimization in the final stage. The final model includes BASS core domain with extended parallel helices at the N-terminal end.

### Engineering *Drosophila* and human mini-tetramer kinesin-5 motors

Human and *Drosophila* kinesin-5 mini-tetramers were designed using the BASS-XL x-ray structure as a template. For the KLP61F KMBX and KMBXt mini-tetramer constructs, the BASS XL was extended by 20 residues based on the heptad pattern observed in BASS-XL the structure (residues 597-833) and fused at its N-terminal end to the KLP61 F motor and neck linker domain (residues 1-369) and were either contained or lacked a C-terminal extension of the KLP61F tail domain (residues 910-1066) with a C-terminal his tag. For the Eg5 mini-tetramer hMBX and hMBXt constructs the *Homo sapiens* Eg5 motor domain and neck linker (1-374) were fused to the N-terminal end of the *Drosophila melanogaster* BASS-XL domain (597-799), either contained or lacked the *Homo sapiens* Eg5 C-terminal tail domain (913-1056), and a C-terminal 6x-His tag with mutations (C25V, C43S, C87A, C99A, N358C, C964S, and C1003S) to allow for the specific labeling of the motor at a single reactive cysteine residue in the neck linker (N358C). Tubulin was purified from Pork or Bovine brains (Castoldi and Popov, 2003). After purification, tubulin was cycled or labeled (with Alexa-546 or Alexa 647 (Thermo Fisher Scientific; Waltham MA), HiLyte 488 (AnaSpec; Fremont CA) or Biotin-LC-NHS (Thermo Fisher Scientific)) and then cycled prior to use. Unless otherwise stated, all chemicals and proteins were purchased from MilliporeSigma (Burlington, MA).

### Motility assays for Drosophila Kinesin-5 (kMBX and kMBXt) minitetramers

Flow chambers were assembled from N 1.5 glass coverslips (0.16 mm thick; Ted Pella) that were cleaned with the Piranha protocol and functionalized with 2 mg/mL PEG-2000-silane containing 2 μg/mL biotin-PEG-3400-silane (Laysan Bio) suspended in 80% at pH 1 (Henty-Ridilla et al., 2016). After the flow chamber was assembled, 0.1 mg/mL NeutrAvidin (Thermofisher) was used to functionalize surfaces. Biotin and Alexa-Fluor-633-labeled porcine tubulin were generated in the laboratory as described (Al-Bassam, 2014) and were polymerized using the non-hydrolysable GTP analog guanosine-5’-[(α,β)-methyleno] triphosphate (termed GMPCPP; Jena Biosciences) or using the MT stabilizing drug, Paclitaxel (sigma). These MTs (100-200 μg/mL in BRB-80: 80 mM PIPES, 1 mM MgCl_2_ and 1 mM ETGA; pH 6.8, 1% glycerol, 0.5% pluronic-F127, 0.3 mg/ml casein, 3 mM BME, 4 mM ATP-MgCl_2_) were flowed into chambers and attached to glass via biotin-neutravidin linkage. Flow chambers were then extensively washed with imaging buffer (25 mM HEPES, 25-150 mM KCl, pH 7.5, 10 mM beta-mercatopethanol; 1% glycerol, 0.5% Pluronic-F127, 0.3 mg/ml casein, 3 mM BME, 4 mM ATP-MgCl_2_). Kinesin-5 MT-stimulated motility was reconstituted at 25°C by injecting 1-20 nM FL-Eg5-GFP combined with a photo-bleach-correction mixture into flow chambers (Telley et al., 2011). Movies were captured in TIRF mode using a Nikon Eclipse Ti microscope using 1.5 Na objective and an Andor IXon3 EM-CCD operating with three (488 nm, 560 nm and 640 nm) emission filters using alternating filter wheel in 2 s increments operated using elements software (Nikon).

### Motility assays for human Kinesin-5 (hMBX and hMBXt) mini-tetramers

Coverslips (22×30mm; Thermo Fisher Scientific) were cleaned by soaking in acetone for 10 minutes, sonicating in 50% methanol for 20 minutes, sonicating in 0.5M KOH for 20 minutes, and then rinsing in MilliQ water three times before drying using nitrogen gas. Cleaned coverslips were stored covered at room temperature. Immediately prior to silanization, coverslips were plasma treated on “high” (~18W; 200mTorr) for 45 seconds after evacuating the chamber. Subsequently, they were soaked in PlusOne Repel-Silane ES (GE Healthcare; Chicago IL) for 20 minutes, transferred to 95% ethanol for 5 minutes, and then sonicated in fresh 95% ethanol for 10 minutes. Finally, they were dried again using nitrogen gas and stored covered at room temperature for up to 2 months.

Taxol-stabilized MTs were prepared as follows: purified unlabeled, Alexa 647-labelled, and biotinylated tubulin were combined at 50μM in BRB80 in a 50:2:1.5 ratio and supplemented with 2mM GTP. After incubating at 37°C for 40 minutes, 40μM Taxol (Cytoskeleton; Denver CO) was added, and MTs were incubated for an additional 30 minutes. Subsequently, MTs were pelleted by centrifugation (8000 rcf, 10 minutes, 25°C), and, after washing the pellet, resuspended in fresh BRB80 supplemented with 40μM Taxol. The pelleting, washing, and resuspension were repeated twice sequentially.

Single molecule motility assays were carried out as follows: Flow chambers with a volume of ~20μL were prepared from a silanized coverslip, a glass slide, and double-sided tape. NeutraAvidin was introduced at 20μg/mL and incubated for 5 minutes, before blocking the remaining surface using 50mg/mL Pluronic F-127 for 30 minutes. Stabilized MTs containing biotinylated tubulin were then flowed in at a concentration of ~0.14mg/mL and allowed to bind for 5 minutes, at which point any unbound MTs were removed by washing the chamber with BRB80 supplemented with 20μM Taxol. Just prior to imaging, the assay buffer consisting of BRB80 supplemented with 20μM Taxol, 0.5mg/mL BSA, 10mM DTT, 2mM ATP, 0.5mg/mL glucose oxidase, 7mg/mL glucose, 0.2mg/mL catalase, 0.2%w/v PEG, 40mM potassium acetate, and ~22nM kinesin-5 (with or without the tail) was introduced. The flow chamber was then sealed at either end using vacuum grease and imaged immediately for no more than 30 minutes. Assays with AMP-PNP were performed identically, except that motors were diluted 36x more and the ATP was substituted with 2mM AMP-PNP (Roche Diagnostics; Indianapolis IN).

TIRF microscopy imaging was performed using an Eclipse Ti-E inverted microscope (Nikon; Melville NY) equipped with diode lasers (100mW; 405nm, 488nm, 561nm, and 640nm) (Coherent; Santa Clara CA), custom optics for TIRF, a 1.49 NA 100x objective, and an additional 1.5X lens to increase the magnification. Two-color image series were acquired by capturing images of the MTs (640nm, 100ms exposure, 1mW) and the motors (561nm, 200ms exposure, 2mW) in an alternating fashion by alternating the laser illumination in synchrony with the rotating filter wheel using a custom-written LabView program, resulting in a frame rate of ~1.2s^-1^. Images were captured on an iXon U897 EMCCD camera (Andor Technology; South Windsor CT).

### Single Molecule Analyses for *Drosophila* kinesin-5 (kMBX and kMBXt)

Motility, run length and run time analyses were carried out as follows: Image movie stacks were pre-processed with photobleach correction and image stabilization plugins using the program FIJI (Schindelin et al., 2012). For motility along individual MTs, individual kMBX and kMBXt motor motility events were identified along anchored MTs based on kymographs in generated for multiple channels. The FIJI kymograph TrackMate plugins (Schindelin et al., 2012) were used to measure particle motility rates, identify their run lengths as and run time. Large collections of motile events along fields of MTs were collected for kMBXt motors at 25, 50, and 100, 125 and 150 mM KCl conditions and collected for kMBX motors at 50 and 100 mM KCl conditions (Table 2). Average MT parameters were determined by frequency binning the motility events in a range conditions and then fitting these events using Gaussian distributions using the program Prism (Table 2). In general, all parameters fit single Gaussian distributions. Run lengths were fitted using exponential decay to identify the half-length for each motor condition. T-tests were performed to determine significance of the differences observed.

The stoichiometry of kMBX and kMBXt motors per multi-motor cluster were determined as previously described (Pandey et al., 2021). Briefly, following correction for the uneven illumination of images and background subtraction, intensities of all NG-labeled kMBX or kMBXt in the first frame of a time-lapse sequence were measured using the TrackMate plugin of the ImageJ-Fiji software(Schindelin et al., 2012; Tinevez et al., 2017). Since kMBX and kMBXt are homo-tetrameric, each motor contains at least four NGs per tetrameric motor. The major peak of the intensity distribution histogram of these mNG-labeled mini-tetramers was fitted to a Gaussian distribution. The center of the Gaussian peak lay at ~600 a.u., which corresponds to the average intensity of single kMBX or kMBXt motor containing one, two, three, or four fluorescent mNG, with each fluorescent mNG molecule contributing ~240 a.u. to the total intensity. Thus, the neon green labeled motor population within this Gaussian peak likely represents single kMBX or kMBXt motors. By this method, we assigned intensity ranges for kMBX or kMBXt molecules fluorescence as <960, 960-1920 and >1920 for single mini-tetramer, pairs mini-tetramers, and higher order oligomers of mini-tetramers, respectively. All the fluorescence intensity measurements to assign cluster size to an kMBX or kMBXt molecule were performed only in the first frame for each data set, thereby significantly reducing the possibility of photobleaching effects.

### Single molecule analyses of Human kinesin-5 (hMBX and hMBXt) mini-tetramers

Image series were drift corrected using the MT images with the Image Stabilizer plug-in for ImageJ. The output coefficients were then applied to the motor images using the Image Stabilizer Log Applier plug-in. These transformation coefficients were saved and used for later steps in the analysis. For images presented in the text, the MT channel was bleach corrected using the Histogram Matching algorithm in ImageJ.

Single molecule tracking was performed on the original, unprocessed images using the TrackMate (Schindelin et al., 2012; Tinevez et al., 2017) plug-in for ImageJ. Briefly, sub-pixel localization was performed using a Laplacian of Gaussian (LoG) filter with an estimated spot diameter of 0.4μm to detect motors. Spots detected in sequential frames were then linked using the Linear Assignment Problem (LAP) tracker using a maximum displacement of 0.6μm, allowing for a two-frame absence of spots, and not permitting track merging or splitting. No filtering of the detected spots or tracks was done in ImageJ.

Quantitative analysis of motor motility activities was performed as follows: MTs were manually traced out in ImageJ, and the x, y coordinates of these ROIs were saved and imported into MATLAB. Detected trajectories >3 frames were then projected along the axis of their respective MT for run length and velocity analysis. Trajectories with a total run length <150nm were not included the analysis to filter out statically-bound motors (e.g., those adsorbed on the coverslip near a MT). All trajectories within 600nm of another MT, or within 600nm of either end of the MT, were ignored in the analysis of pausing/motility. The local α-value analysis was performed on all trajectories with >10 frames using drift-corrected (but not projected) x, y localizations. These were linearly interpolated to account for the slightly varying frame rate. α-values were calculated in sequential 8-frame windows using delays of 2, 3, and 4 frames. Change points were detected in the resulting α-values using the findchangepts function in MATLAB (The MathWorks; Natick MA) with a minimum improvement in the residual error of 0.3 and a minimum distance between consecutive change points of 3 frames. The means between detected change points were then evaluated and sections with a mean α-value >1.1 were deemed processive. The localizations in the original trajectory were then assigned as processive or paused based on the classification of the interpolated frame that was nearest to them in time. Subsequent analysis of intensities, velocities, and inter-pause run lengths was performed using these classifications. Fitting was done using maximum likelihood estimation (MLE). For the velocity histograms, normal distributions were fit using the central 96% of the data. For pause time estimates, distributions were fit using the lower 95% of the data because of the presence of a small number of inactive motors that are classified to have long “pauses”. For analysis of trajectories in the presence of AMP-PNP, no minimum run length was used, and the α-value histogram was fit to the upper 95% of the data to ignore the skew.

MT plus-End motor localization intensity analyses were performed as follows: The duration of time motors/clusters were present at the plus-ends of MTs was quantified using ImageJ. Briefly, all unobstructed MT plus-ends (identified based on the direction of motor motility) were marked using point regions of interest (ROIs), which were added to the ROI Manager. These were then converted to circular ROIs with a diameter of 7 pixels centered on the point using a custom-written script. The intensity in these ROIs was traced over time using Multi Measure, ensuring that the recorded measurements included standard deviation, min & max gray value, center of mass, integrated density, and mean gray value. This process was also repeated for 9 background spots spread throughout the field of view. The output from each movie was saved, and all data were imported into MATLAB for further analysis. A moving average was applied to the raw integrated density signal (IDS). In every ROI, a cluster/motor was deemed present in a given frame of the movie if this filtered IDS in the ROI exceeded that of background ROIs by 3 standard deviations. Only cluster lifetimes >9 frames were considered based on the approximate time taken for a motor to walk through the ROI. The lifetimes of any clusters present in the first or last frame of the movie were marked as censored for the subsequent lifetime analysis. Lifetime analysis was performed by fitting the empirical cumulative distribution function (CDF) to a mixture of two Weibull CDFs using MLE, taking censoring into account.

### Reconstitution of full length and mini-tetramer Kinesin-5 MT crosslinking and sliding

MTs for surface-immobilization were generated via mixture of HiLyte 647 tubulin (TL670M), biotinylated tubulin (T333P), and unmodified tubulin (T240) at a ratio of 1:1:20 along with 1mM GMPCPP. MTs were polymerized at 37°C for 1 hour before clarification and stabilization in 30uM Taxol following published protocols (Shimamoto et al, 2015). ‘Free’ MTs for gliding were generated via mixture of rhodamine tubulin (TL590M) and unmodified tubulin at a ratio of 1:20 along with 1mM GMPCPP, and were polymerized, clarified, and stabilized in 30uM Taxol following similar protocols.

The flow chamber design and assay preparation were modified from a previously described protocol (Shimamoto et al., 2015). Anti-parallel MT bundles were constructed using passivated glass coverslips coated with SVA-PEG at a ratio of 50 PEG:1 biotin-PEG. All reagents were prepared with 1X BRB80 buffer. Following each reagent flow-in and incubation, a flush with ~3 chamber volumes of BRB80 was performed. Reagents were introduced stepwise with the following order and incubation times: (1) 0.5 mg/mL neutravidin, 2 minutes; (2) 0.5 mg/mL alpha casein surface block, 3 minutes; (3) HiLyte-647 biotinylated MTs with 0.2mg/mL alpha casein with no additional incubation and immediate flush; (4) kinesin-5 construct at desired final concentration as reported with 0.2 mg/mL alpha casein, 2 minutes; (5) Rhodamine 561 MTs with 0.2 mg/mL alpha casein, 5 minutes, and the corresponding chamber flush included 1 mM TCEP bond breaker solution in BRB80; (6) imaging buffer (25 mM HEPES, 25-150 mM KCl, pH 7.5, 10 mM beta-mercatopethanol; 1% glycerol, 0.5% Pluronic-F127, 0.3 mg/ml casein, 3 mM BME, 4 mM ATP-MgCl_2_) with Oxygen Scavenging System (4.5 mg/ml glucose, 350 U/ml glucose oxidase, 34 U/ml catalase, 1 mM DTT) was then used to flush any unattached MTs. The chamber was then sealed with clear nail polish prior to experiments.

MT bundles were imaged using three-channel TIRF microscopy using the following exposure times and laser lines from a Nikon LUNA four-channel laser launch module: HiLyte-647 MTs: 640 nm laser (60% power, 200 ms exposure); GFP-tagged kinesin-5: 488 nm laser (30% power, 100 ms exposure); and rhodamine MTs: 561 nm laser (30% power, 100 ms exposure). Imaging was performed on a Nikon Ti-E inverted microscope with a CFI Apo 100X/1.49NA oil immersion TIRF objective. Images were acquired using a Photometric Prime 95B camera controlled with Nikon NIS Elements software. Analysis of fluorescent data and generation of intensity linescan data sets were performed using FIJI (ImageJ) tools(Schindelin et al., 2012).

## Figure Legends for figure supplements

**Figure 1 figure supplement 1. Details of the BASS-XL x-ray crystal structure determination.**

A) Purification and Crystallization of the BASS-XL protein: left panel, Size Exclusion Chromatography (SEC) trace showing the purified BASS-XL. Middle panel, SDS-PAGE of purified BASS-XL. Right panel, images of BASS-XL C2 crystals used for x-ray structure determination.

B) Alignment of the BASS (grey) and BASS-XL (red) N-terminal structure ends reveal the extension of the BASS-XL and formation of the heptad repeats at the N-terminal end.

C) 2Fo-Fc map revealing the final model for BASS-XL built using the BASS-XL crystallographic data at 4.4-Å resolution. The model reveals the detailed helical coiled-coil dimers at the BASS-XL N-terminal end.

D) Crystallographic packing organization of BASS-XL minifilaments within the C2 space group for crystals used for structure determination.

**Figure 1 figure supplement 2. Designing kinesin-5 mini-tetramers using the BASS-XL x-ray structure as a short mini-filament template.**

A) Domain organization of designer Drosophila kinesin-5 mini-tetramers. KLP61F motor and neck linker domain (orange: residues 1-369) were fused to an extended BASS-XL minifilament, with its coiled-coil extension (green), dimerized region (cyan), tetrameric region (red) and C-terminal stabilizing zone (blue). The kMBXt construct included fusing the C-terminal kinesin-5 tail domain (residues 910-1036) to the C-terminal domain of the BASS-XL minifilament.

B) SEC-trace and SDS-PAGE for a peak fraction of the bacterially expressed of the C-terminally neon green (mNG) tagged KMBX and kMBXt motors.

C) Top, a side view of a structural model revealing the organization of the designed bipolar mini-tetrameric kinesin-5 motor with various regions (colored as described A) shown. The model reveals the motor neck-linker domains are oriented to face in different directions due to their positioning on the BASS-XL minifilament and the mini-tetrameric motor is 38-nm in length, which is roughly half the length of native kinesin-5. Below, a 90° rotated view of the view shown on top. D) a subunit colored view of the kinesin-5 mini-tetrameric motor design revealing the subunit organization of the motor and minifilament domains. Below, a 90° rotated view of the view shown on top.

**Figure 3-figure supplement 1.**

**A)** Additional traces of parsing analysis for trajectories of hMBXt, similar to Figure 1d. Pauses (red) and periods of processive motility (blue) were identified in trajectories (top plots) based on the local α-value calculated within a sliding window along the trajectory, which fluctuated between ~2 and ~0 (bottom plots).

**B)** Histograms showing the number of distinct sections identified as paused for hMBXt motors (purple) and hMBX motors (green) in a given trajectory as a fraction of the number of long trajectories (n = 143 for hMBXt and n = 111 for hMBX).

**C)** the number of all trajectories >3 frames (n = 1406 for hMBXt motors and n = 2056 for hMBX). This shows that fewer of the trajectories for hMBX motors were long enough to use in our parsing analysis and, as short trajectories were less likely to contain pauses (see, for example Figure 2C), shows that a small fraction of total trajectories contained pauses for hMBX motors.

**D, E)** MSD curves for paused (red) and processive (blue) sections for hMBXt **(D)** and hMBX **(E)** motors plotted on a linear scale, revealing the parabolic shape characteristic of super-diffusive motility for processive sections, and a horizontal line for paused sections. Lines were fit to <x^2^>=(v^2^+σ_v_^2^)τ^2^+2ε^2^ and <x^2^>=2ε^2^, respectively. Error bars represent SEM. For a, n = 143 trajectories from 9 independent experiments. For b, n = 111 trajectories from 3 independent experiments.

**F)** Kymograph showing the static binding of mini-tetramers in the presence of AMP-PNP. Scale bar 10s wide and 1μm tall.

**G)** Repeating the local α-value analysis for mini-tetramers with the tail domain in the presence of AMP-PNP (black; dotted line) reveals a uni-modal distribution, similar to the mean α-value of the paused sections (red; solid line) in the presence of ATP. In contrast, the mean α-value of processive sections (blue; solid line) in the presence of ATP is closer to 2. For AMP-PNP, n = 442 trajectories from 2 independent experiments. For ATP, n = 143 trajectories from 9 independent experiments.

**Figure 4-figure supplement 1**. Measuring kMBXt clustering using tracking and intensity analysis.

A) Low magnification view of the MT (left) and kMBXt (right field) prior to analysis.

B) The intensity analysis (right) for all spots detected (left) identified via TrackMate plugin using FIJI/imageJ. The kMBXt intensity histogram is presented was converted into cluster values based on the fluorescence intensity of NG per motor subunit within each mini-tetramer. Details are described in materials and methods.

**Figure 4 figure supplement 2: Example kymographs for kMBXt clustering at increasing ionic strengths**

Example kymographs of kMBXt clustering along MTs at (A) 25 mM, (B) 50 mM, (C)100 mM, (D) 125 mM KCl and (E) 150 mM KCl conditions. The kymographs show that clustering increases dramatically from 25 to 100 mM KCl but it decreases at 125 and 150 mM KCl. The MT plus-end accumulation seems to concentrate the largest kMBX clusters (left side of each kymograph). (F) Description for types of cluster and motor merging events described in Figure 4 figure supplement 4B.

**Figure 4-figure supplement 3:** Characterizing kMBXt motility properties in relation cluster formation. Distribution for motility velocity A), Run length. B) and Run time. C) for kMBX (green) and kMBXt (purple) at 25-150 mM KCl. The types of clusters formed for kMBXt at different conditions. For each type a smaller intensity spot merges with larger intensity spot (i.e., 1+2). These intensities were delineated using the approach presented in Figure 4 figure supplement 1 and described in materials and methods. D) Relationship of kMBXt motility velocity (left), Run length (middle) and run time(right) to cluster size for the 25 mM (green), 50 mM (pink), 100 mM (Gray), 125 mM (Cyan) and 150 mM KCl (orange). These data show that cluster size does not correlate to any of the properties described.

**Figure 4 figure supplement 4: Analyses of MT plus-end accumulation of kinesin-5 minitetramers**

**A)** Clusters, which generally arrive at plus-ends as assemblies and dissociate as assemblies, were detected based on the intensity in a ROI over time. Blue line denotes intensity trace in ROI at the plus-end of a MT. Magenta spots identify frames in which no motor is identified as present. Green spots identify frames in which motor is identified as present. Other thin lines denote intensity traces of background ROIs used as a reference to identify the presence of a motor. Inset: An example of a motor arriving and falling off a plus-end is shown. 20 frames between time points. Scale bar 1μm wide.

**B)** Histograms of the frame-to-frame velocities for single motors (light purple; dotted line) and clusters (dark purple; solid line). During the processive sections of trajectories, clusters move more slowly than single motors. Their velocities during paused sections are similar. For a and e, n = 143 trajectories from 9 independent experiments for hMBXt and n = 111 trajectories from 3 independent experiments for hMBX. For b and c, n = 351 plus-end clusters detected from 8 independent experiments for hMBXt motors and n = 211 plus-end clusters detected from 3 independent experiments for hMBX motors.

C) Additional traces of cluster detection at the plus-ends of MTs, similar to Figure 4C. Clusters generally arrive at plus-ends as a whole and dissociate as a whole, regardless of the plateau intensity, but we did observe single motors appearing to join a pre-existing cluster, as well as some clusters forming at the plus-end (e.g., top left). Blue line denotes intensity trace in ROI at the plus-end of a MT. Magenta spots identify frames in which no motor is identified as present. Green spots identify frames in which motor is identified as present. Other thin lines denote intensity traces of background ROIs used as a reference to identify the presence of a motor. Top four traces for hMBXt motors and lower two for hMBX motors.

D) The cumulative distribution function for the mixed Weibull distribution used to fit the empirical cumulative distribution of lifetimes for hMBX and hMBXt motors, as well as the obtained estimates of the parameters. Note that the short lifetimes and shape parameters are similar, while the longer lifetime is longer for hMBXt motors. Notably, the weighting of the shorter lifetime is also lower for hMBXt motors, suggesting that a larger fraction of motors is in the longer lifetime population.

E) An example of a cluster of human kinesin-5 HMBXt motors dragging the plus-end at which the motors accumulated along another MT, acting to align the two MTs. This demonstrates that the plus-end clusters are still motile and are able to withstand load (the MTs are quite bent and still anchored to the coverslip at some sites by an avidin-biotin interaction). Note that to observe this, a higher density of MTs was immobilized using a lower density of Neutravidin. 50 frames between time points. Scale bar 2μm wide.

**Figure 5 figure supplement 1:** Analyses of **Velocity of MT sliding for native and minitetramer kiensin-5:** Individual frames from time-lapse image series were processed via linescan analysis to determine MT minus-end positions. These values are then plotted as a function of time for (a) hFLt and (b) kMBXt proteins. Velocities are calculated by applying a sliding window function of width 5 frames to the position data and a linear slope is calculated for the corresponding data sets (b,d). Velocity values below 15 nm/s for more than 3 consecutive frames were then classified as pausing events (red dashed line in b,d).

## Video Legends

**Video 1:** Large-scale view of hMBXt motors (green) undergoing motility along MTs (magenta) revealing kinesin-5 mini-tetramer motor pausing, clustering, and MT plus end association.

**Video 2:** Close-view of hMBX motors (green) undergoing motility, forming clusters, with pausing and while dwelling at MT (magenta) plus-ends.

**Video 3:** motility of kMBXt motors at four different ionic strength conditions (25-150 mM KCl) revealing the impact of ionic strength on motor clustering, processive motility and plus-end attachment. Clustering is improved at 50 mM KCl and decreases at 100-150 mM KCl. MT plus end accumulation occurs at 25-100 mM KCl and is diminished at 150 mM KCl.

**Video 4:** An example of a cluster of kinesin-5 hMBXt motors dragging the plus-end at which the motors accumulated along another MT, acting to align the two MTs. This demonstrates that the plus-end clusters are still motile and are able to withstand load (the MTs are quite bent and still anchored to the coverslip at some sites by an avidin-biotin interaction). Note that to observe this, a higher density of MTs was immobilized using a lower density of Neutravidin. 50 frames between time points. Scale bar 2μm wide.

**Video 5:** Three examples of MT sliding by native kinesin-5 (hFLt). The hFLt motors (green) slide a paired MT (red) along the anchored MT (magenta).

**Video 6:** Three examples of MT sliding by mini-tetramer-kinesin-5 (kMBXt). The kMBXt motors (green) slide a paired MT (red) along the anchored MT (magenta).

## Notes

### Competing Interest Statement

The authors have declared no competing interest.

