## Supplementary material for "The mechanochemical origins of the microtubule sliding motility within the kinesin-5 domain organization": Figure 1-5, Figure supplements

Figure 1-sup 1

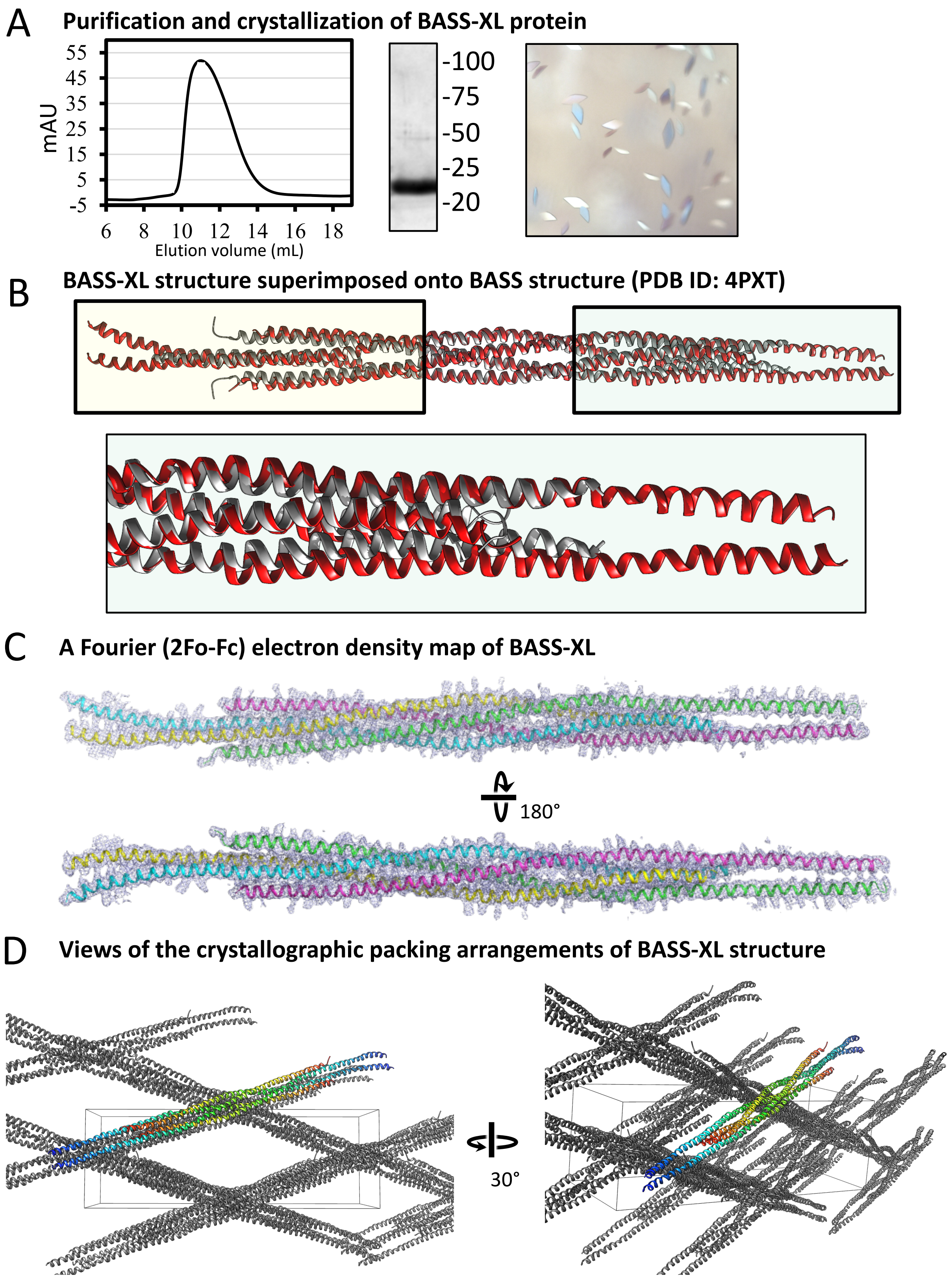

Figure 1-sup 2

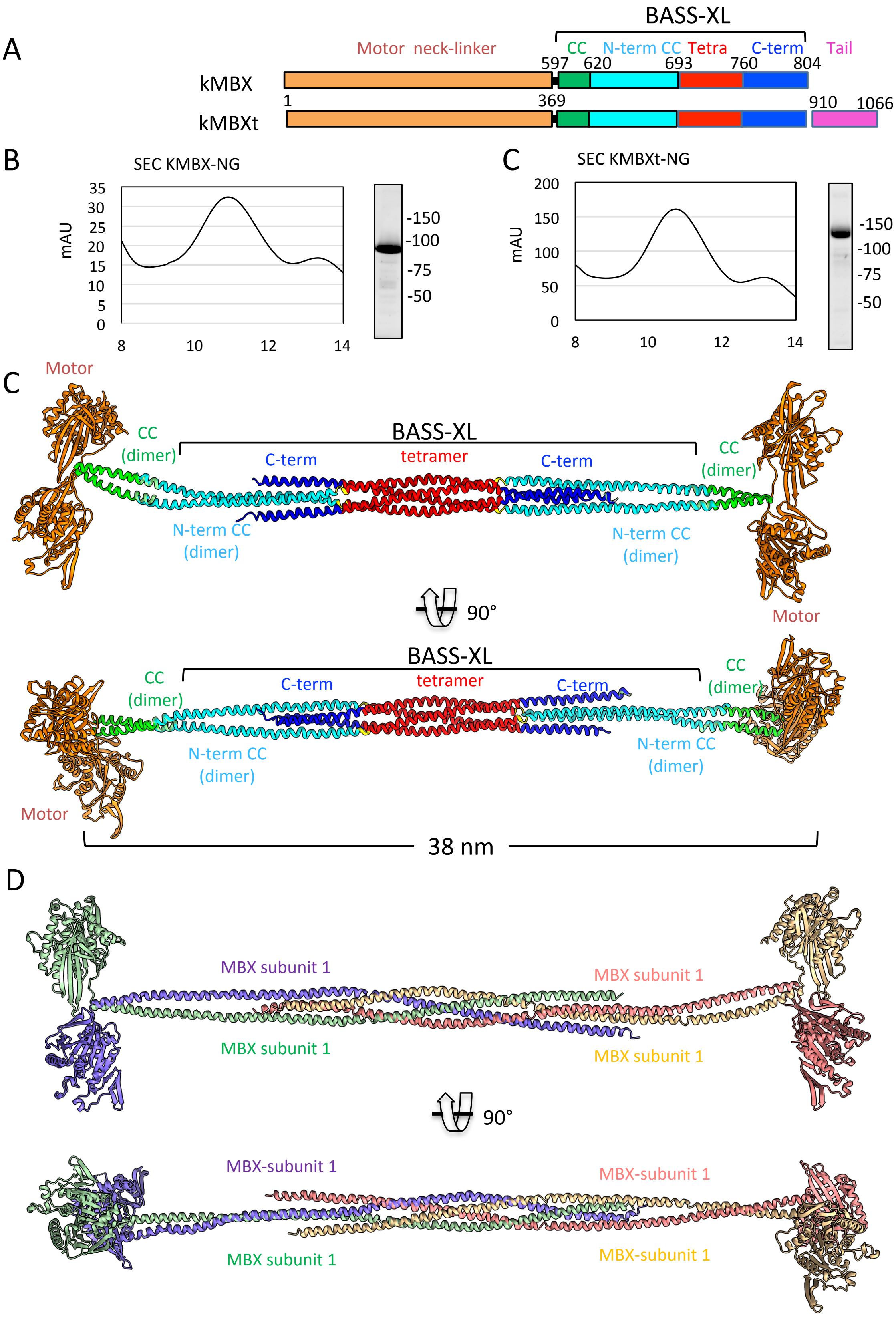

Figure 3-sup 1

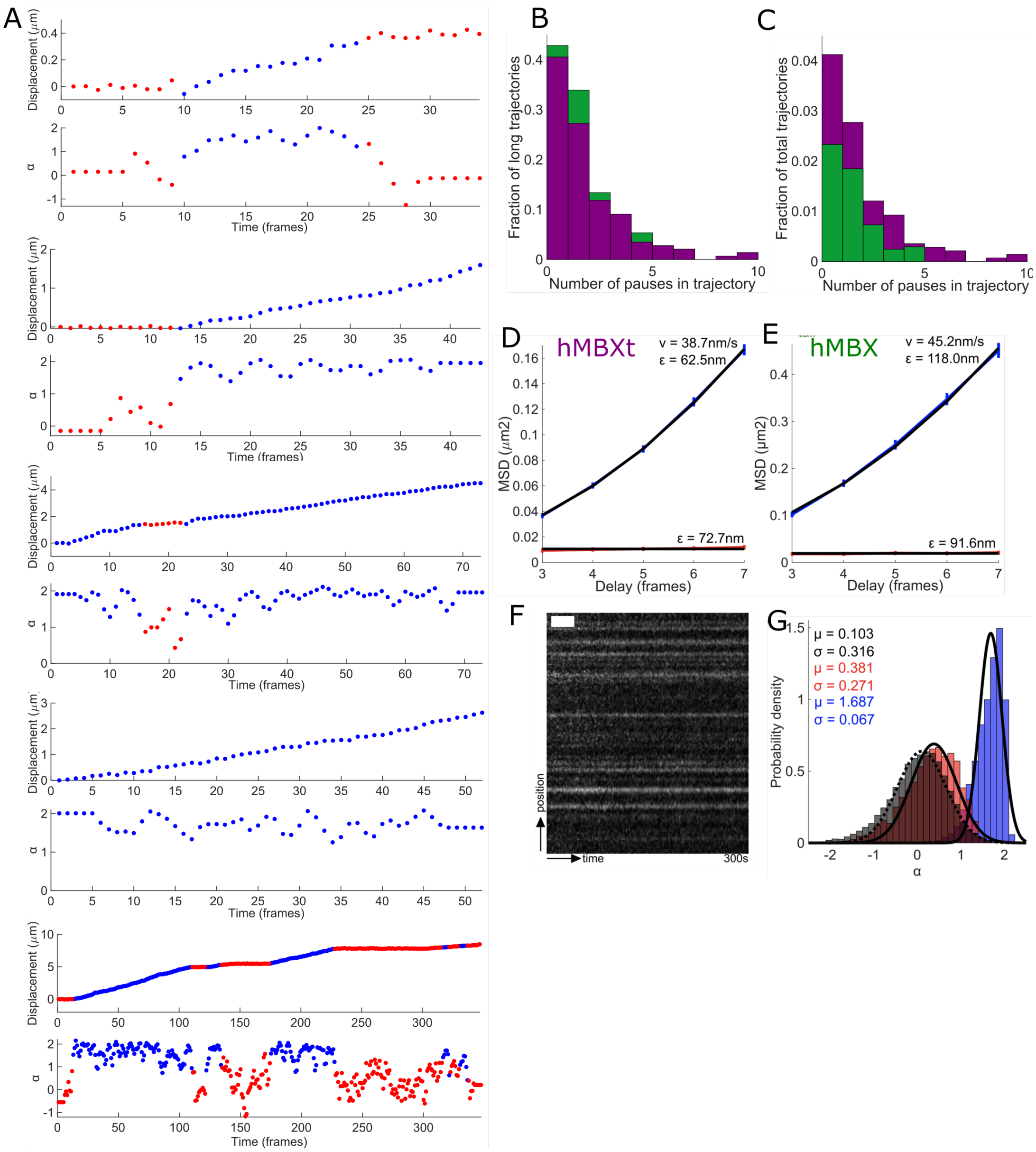

Figure 4-sup 1

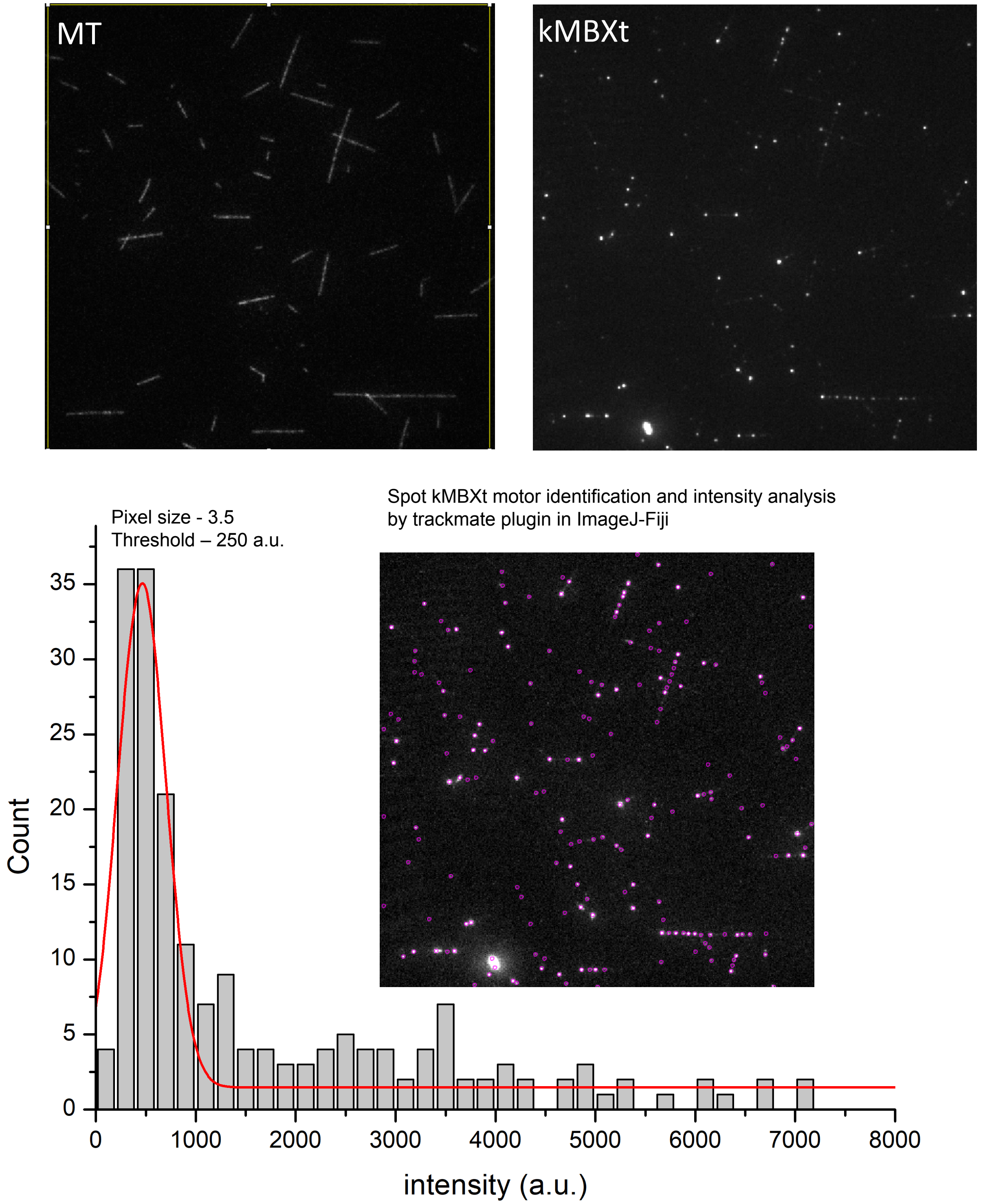

Figure 4-sup 2

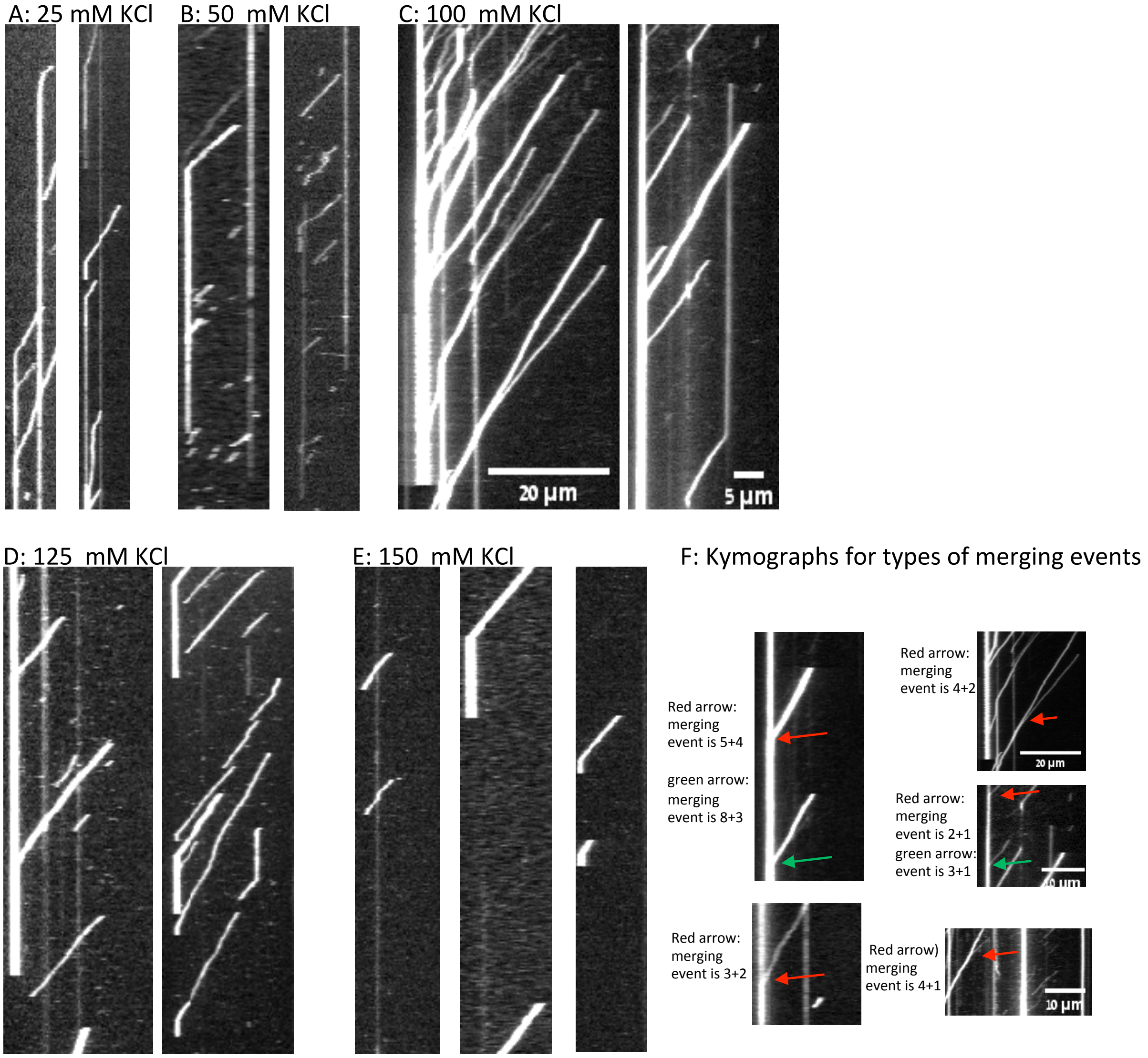

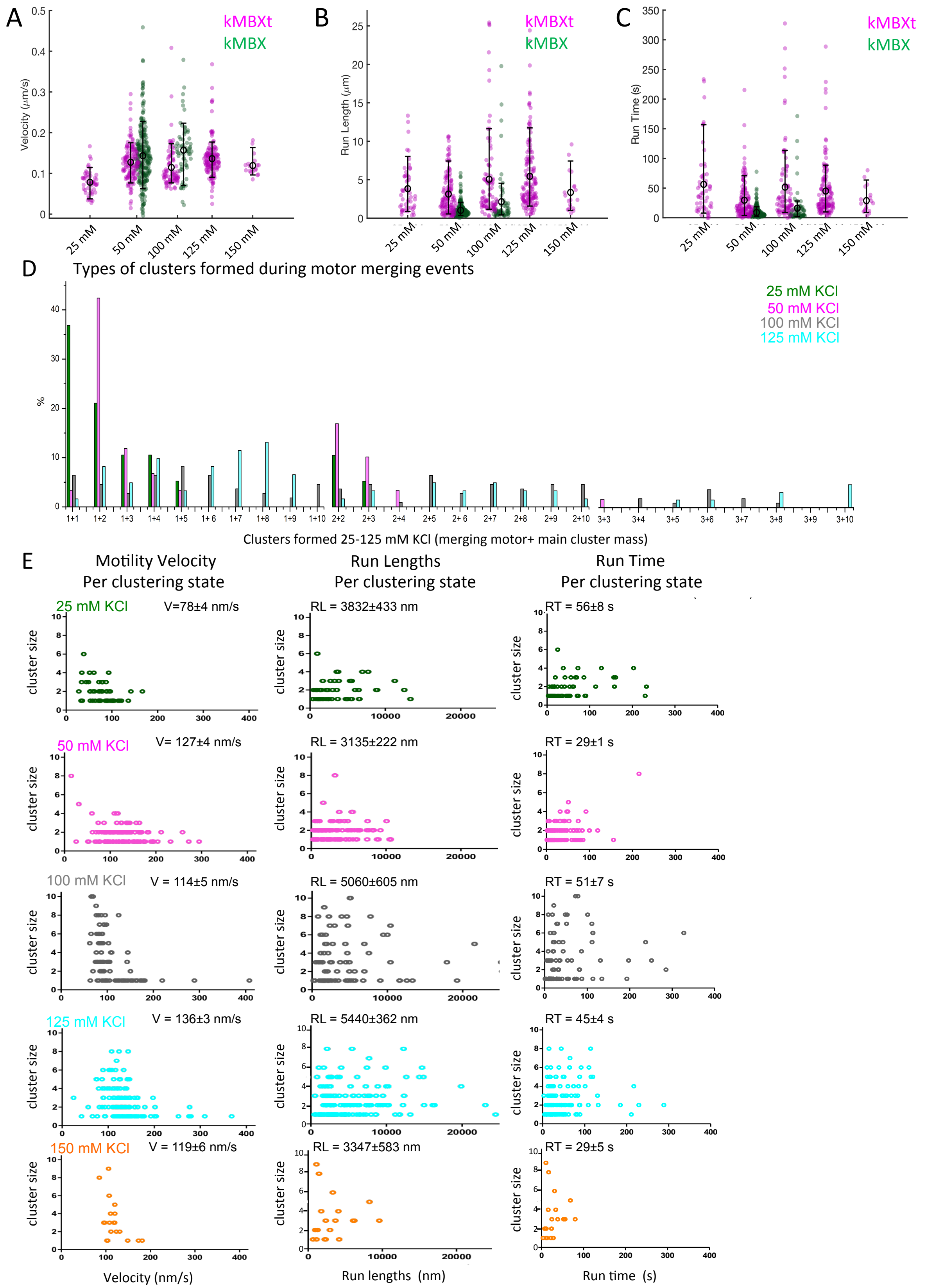

A
kMBXt
kMBX
Velocity (μm/s)
25 mM
50 mM
100 mM
125 mM
150 mM
B
Run Length (μm)
C
Run Time (s)
D
Types of clusters formed during motor merging events
25 mM KCl
50 mM KCl
100 mM KCl
125 mM KCl
%
Clusters formed 25-125 mM KCl (merging motor+ main cluster mass)
E
Motility Velocity
Per clustering state
Run Lengths
Per clustering state
Run Time
Per clustering state
25 mM KCl
V=78±4 nm/s
RL = 3832±433 nm
RT = 56±8 s
50 mM KCl
V= 127±4 nm/s
RL = 3135±222 nm
RT = 29±1 s
100 mM KCl
V = 114±5 nm/s
RL = 5060±605 nm
RT = 51±7 s
125 mM KCl
V = 136±3 nm/s
RL = 5440±362 nm
RT = 45±4 s
150 mM KCl
V = 119±6 nm/s
RL = 3347±583 nm
RT = 29±5 s
cluster size
Velocity (nm/s)
Run lengths (nm)
Run time (s)

Figure 4-Sup 4

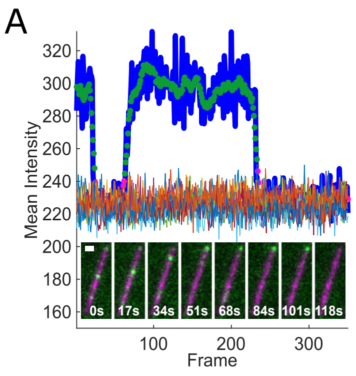


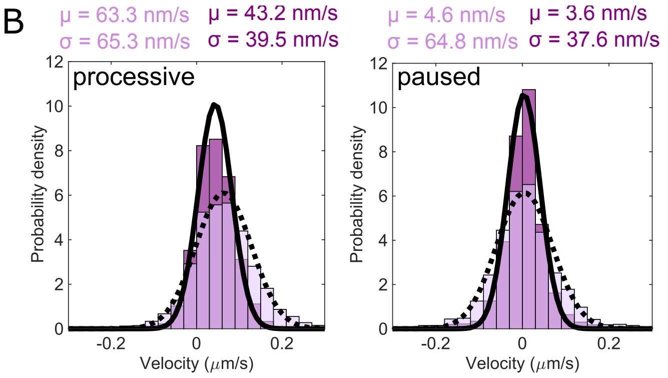


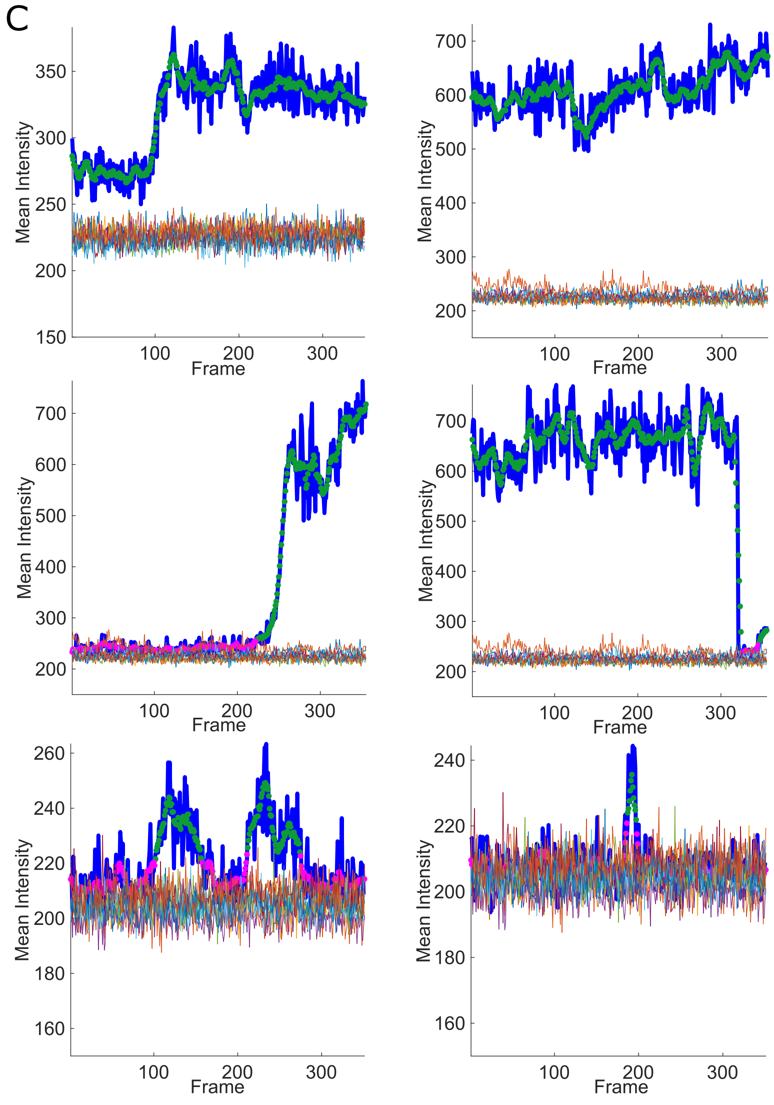


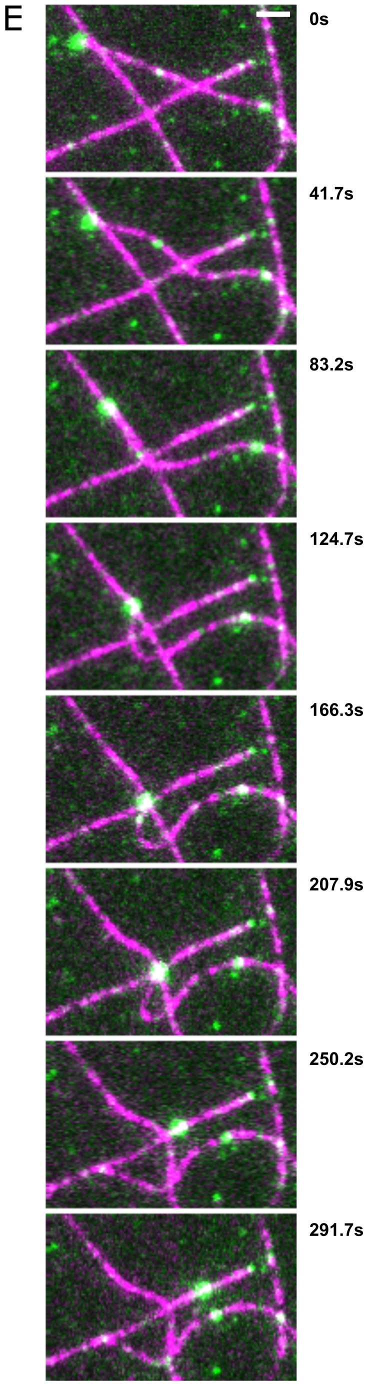


D

$$F(x|a_1, b_1, a_2, b_2, p) = p\left\{1 - e^{-\left(\frac{x}{a_1}\right)^{b_1}}\right\} + (1-p)\left\{1 - e^{-\left(\frac{x}{a_2}\right)^{b_2}}\right\}$$

| | $a_1$ | $b_1$ | $a_2$ | $b_2$ | p |
|---|---|---|---|---|---|
| | scale parameter; lifetime (s) | shape parameter (b<1 if failure rate decreases with time) | scale parameter; lifetime (s) | shape parameter (b<1 if failure rate decreases with time) | weighting of shorter lifetime fit |
| - tail | 14.6 ± 3.2 | 1.07 ± 0.20 | 139.9 ± 137.5 | 0.76 ± 0.42 | 0.74 ± 0.18 |
| + tail | 16.4 ± 4.9 | 0.88 ± 0.12 | 483.7 ± 153.2 | 0.89 ± 0.64 | 0.49 ± 0.15 |

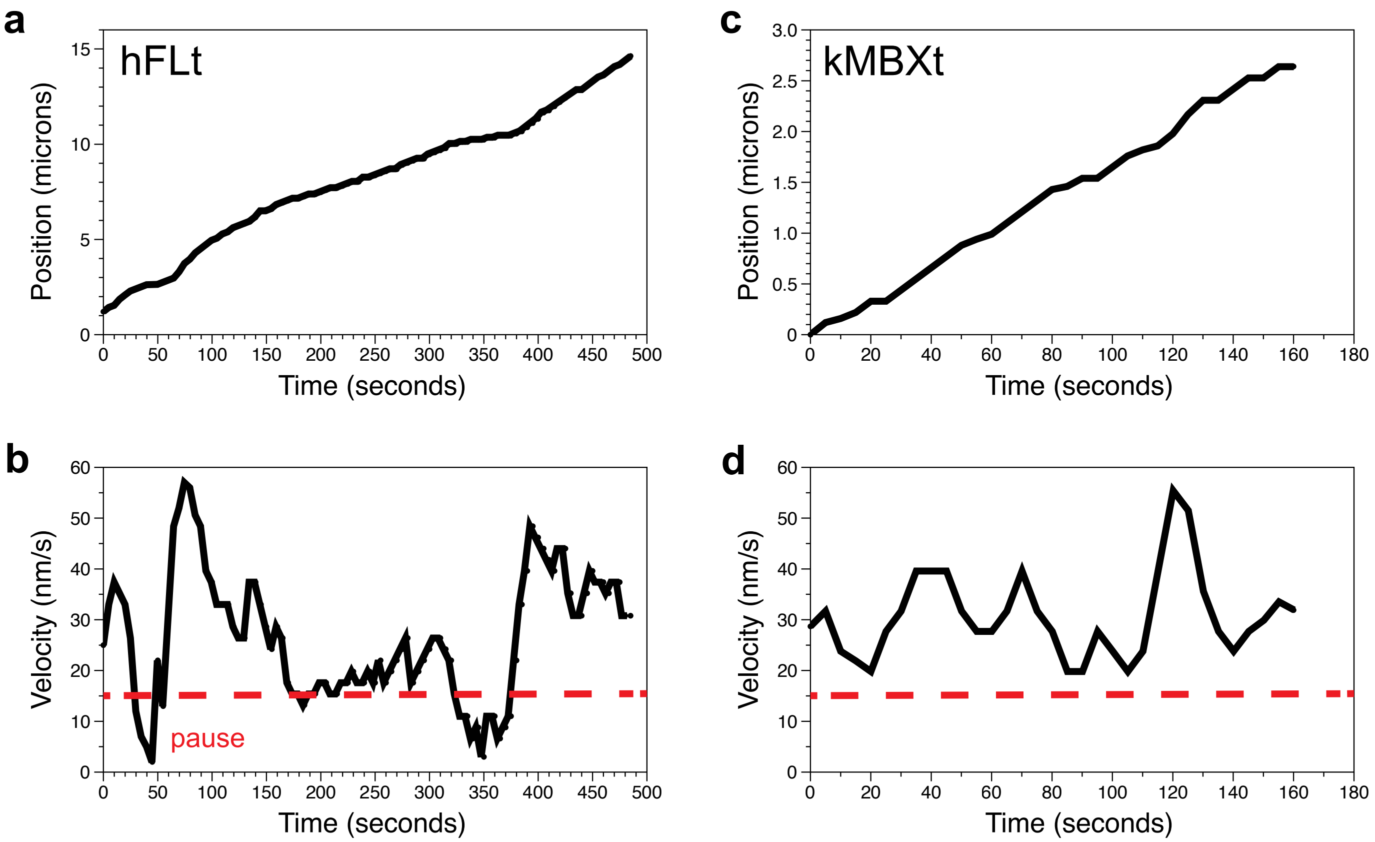
a
hFLt
Position (microns)
Time (seconds)
0
5
10
15
0
50
100
150
200
250
300
350
400
450
500
c
kMBXt
Position (microns)
Time (seconds)
0
0.5
1.0
1.5
2.0
2.5
3.0
0
20
40
60
80
100
120
140
160
180
b
Velocity (nm/s)
Time (seconds)
pause
0
10
20
30
40
50
60
0
50
100
150
200
250
300
350
400
450
500
d
Velocity (nm/s)
Time (seconds)
0
10
20
30
40
50
60
0
20
40
60
80
100
120
140
160
180
